# Phylogenetic reconstructions of Polynesian medicinal plant use reveal adaptive strategies to meet health needs

**DOI:** 10.1101/2022.10.21.513211

**Authors:** Jamie Thompson, Fiona Jordan, Julie Hawkins

## Abstract

Modern migrants using plants to meet their health needs are known to conserve traditional knowledge, but also to innovate to adapt to their new environment. The voyage into Polynesia is amongst the most remarkable of human migrations, resulting in the peopling of isolated, difficult to reach archipelagos. We use this context to determine the role for adaptation in plant-based healthcare at pre-historic timescales. Testing the extent to which the new floristic environments encountered, cultural ancestry or geographic proximity predict the composition of ethnopharmacopoeias, we reveal adaptation to new floristic environments across seven oceanic ethnolinguistic groups. Reconstructions using data that encompass therapeutic applications and plant parts reveal more than three quarters of the plants used cross-culturally have use likely to be innovations by the first migrants into Oceania. For the other plants, there are non-tree-like patterns in therapeutic applications and plant parts used, showing that even when plants have continuity of use there is lability in how they are used. Applying linguistic criteria to the plants with putatively deep cultural uses, we find two, qaoa (*Ficus*) and walo-walo (*Premna*), that are highly conserved in therapeutic use, plant part used and with cognate names. Our study highlights the remarkable flexibility of Polynesian people seeking to meet health needs.

## Main

Changing health needs and availability of plants, acculturation, and increasing access to biomedical pharmaceutical drugs drive changes in knowledge of medicinal plants (1). Innovation and loss of knowledge contribute to a constantly changing ethnopharmacopoeia (2), and loss of knowledge may itself be considered an adaptive community response to changing medical needs (3). Yet documenting continuity and change in medicinal plant use is challenging (3). Studies returning to sites for re-inventory of medicinal plant use are rare, and focussed on rapid erosion of knowledge (4), a phenomenon more often revealed by comparing knowledge held by informants of different ages (5, 6). Diachronic analysis comparing historical materia medica with more contemporary survey data can document transmission or adaptation of knowledge at the timescale of written records (7) though the texts themselves shape the use of the plant-based medicines (8). Recognising that these methods are limited to certain scenarios, phylogenetic comparative methods (PCMs) were recently suggested as a general means to ascertain whether there is continuity of knowledge at the timescale at which ethnolinguistic groups diverge (9).

Functional and group-signifying cultural knowledge is often passed vertically, from ancestor to descendant (10). Though knowledge transmission can be observed between individuals, PCMs are able to explore vertical evolutionary processes occurring over scales that cannot be observed directly. One challenge when seeking to identify putatively deep, ancestral knowledge of medicinal plants is that culturally unrelated groups use similar plants through apparent independent innovation (11). Cross-cultural transmission, or horizontal transfer of knowledge, may also hinder attempts to identify ancestral knowledge. One application of the phylogenetic approach teased apart the contribution of innovation and cross-cultural transmission, versus retention of ancestral knowledge in Nepal. Adaptation to floristic environment was the only significant factor correlated with the overall composition of the ethnoflora (12). The task of understanding the drivers of overall similarity in ethnofloras is separate, but related to the reconstruction of uses of specific plant species in the past, an approach that can identify ancestral knowledge even if plant use is predominantly driven by plant availability. A recent study assessed the possibility of reconstructing ancestral knowledge of medicinal plants in Nordic cultures, evaluating inferences against historical and archaeobotanical sources (13). For many plants, ancestral reconstructions of Old Norse knowledge were supported by historical knowledge, but some were revealed by PCMs alone. There is insufficient archaeobotanical evidence of medicinal plant use in Polynesia to validate our studies in this way. The Norse study provides proof of concept of the phylogenetic methods that we apply here to a situation where other data are sparse. A comprehensive study of continuity and change in ethnofloras would consider both the overall composition of the flora and ancestral reconstructions of its components. The latter depends on reconstructing ancestral knowledge about the plants used, therapeutic applications, parts used and local names (9). These aspects of plant use provide tests of ancestral knowledge and complement investigations at the level of the overall ethnoflora.

Migration is one driver of adaptation in medicinal plant use, since it is a special case of exposure to new floristic environment (14). Following migration, continuity in plant use could be maintained if the preferred species is in the new environment, or if it can be introduced. Otherwise, plants related to, or otherwise similar to, the original preferred species might be adopted (15). As well as encountering new floras, migrant peoples may encounter other cultural groups who hold different knowledge, and cross-cultural transmission between indigenous residents and migrants has been shown to be a source of change in ethnofloras (16). These factors are limited where migration is to previously uninhabited islands (such as Remote Oceania), and when post-settlement cross-cultural transmission is attenuated. Austronesian expansion into the Pacific began 4750 to 5800 years B.P., most probably from Taiwan (17). Within 200 years of the settlement of Aotearoa (New Zealand) and Hawaii, archaeological evidence suggests cross-cultural transmission had declined to negligible levels (18). Austronesian cultures therefore provide a natural experiment for testing the relative importance of adaptation on encountering new plants versus retention of ancestral knowledge in the context of limited opportunity for horizontal transmission at the population level.

We were able to identify seven ethnolinguistic populations/archipelagos for which there was floristic data and adequate data describing plant use. They were Chamorro (Micronesian), Rotuman (Melanesian) and five Polynesian cultures, Hawaiian, Maori, Marquesan, Samoan and Tongan. The Polynesian selection reflects geographically widely dispersed cultures with well-known relationships (Figure 1). For all of these populations it was possible to measure the overall relatedness of the ethnofloras and the floras using community phylogenetic methods. Ethnofloristic distance was the response matrix and floristic distance one of the explanatory matrices for multiple regression on distance matrices (MRM). We incorporated floristic distance in order to estimate the extent to which floristic environment shapes medicinal plant use. A second explanatory matrix was ethnolinguistic distance, a measure of shared ancestry. Geographic distance was incorporated as a proxy for opportunity for cross-cultural transmission, recognising that other data not so readily available might be better indicators of contact in the context of Austronesia (18, 19).

**Figure 1.**
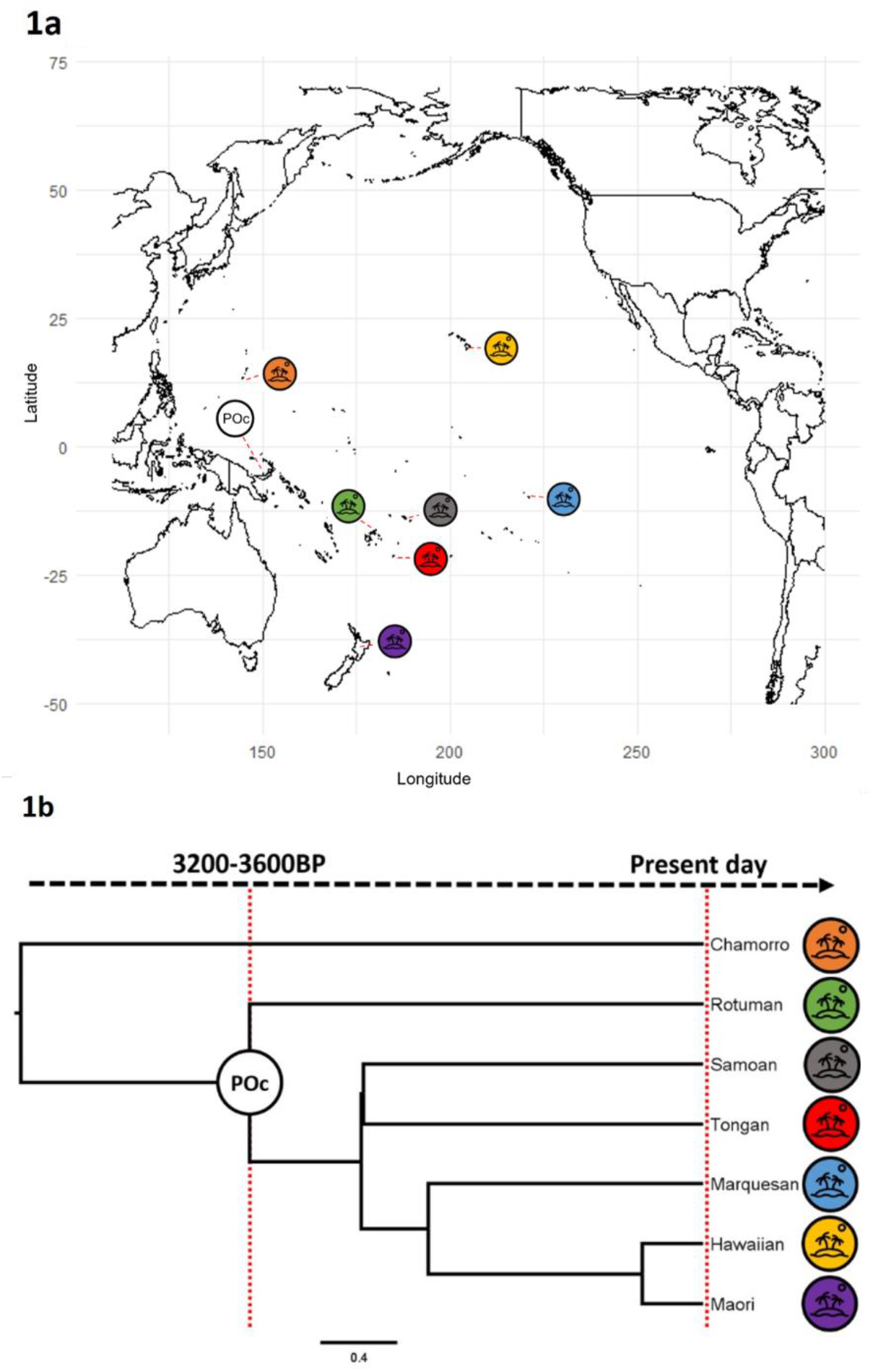
Spatial distribution and linguistic relationships of the ethnolinguistic groups studied. 1a map showing the spatial sampling of groups, with the presumed position of Proto-Oceanic in the Bismarck Archipelagof. 1b language phylogeny showing their relationships. The POc node represents the (putative) Proto-Oceanic ethnolinguistic group. Spatial and linguistic proximities are strongly uncoupled.

We revealed a predominant pattern of adaptation to new floristic environments across the seven oceanic ethnolinguistic groups. All three variables could independently explain the variation in medicinal plants used (R^2^ 0.85, p<0.05). The floristic environment explained the largest proportion of the variance in medicinal plant use, with cultural groups using more similar plants in more similar floristic environments (R^2^ 0.85, p<0.05). Shared ancestry explains approximately 60 times less of the variance than floristic environment (R^2^ 0.015, p<0.05). Geographical proximity contributes the least to similarity in medicinal plant selection (0.0000061, p<0.05). The floristic environment was also the primary driver of plant selection ethnomedicine in the previous study of Nepal (12). However, in the case of Nepal, neither ancestry nor proximity were significantly correlated with composition of the ethnoflora. The present study uses MRM methods rather than Mantel methods and also uses a time-calibrated Bayesian linguistic phylogeny, rather than the estimate of relatedness derived from the Ethnnologue. Nevertheless, this present study, the second using community phylogenetics and matrix methods, contributes to the emerging picture of adaptation to the floristic environment as the predominant process. In changed environments, adaptation is expected to occur rapidly when an aspect of culture is functional. For example, functional aspects of Polynesian canoe design appear most decoupled from the background of shared linguistic ancestry than decorative/symbolic aspects, implying rapid evolution in function (20). Knowledge of plants is associated with health outcomes and this might explain the apparently high rate of adaptation in knowledge of medicinal plants (21, 22). Our study specifically excludes recently introduced plants, but the prevalence of introduced plants described elsewhere also points to adaptive ethnopharmacopoeias (23).

The narrative of rapid, adaptive change in medicinal plant use might lead us to expect change to also be mediated by cross-cultural, horizontal transfer of knowledge, even if there was limited contact. However, although vertical conservation of knowledge and horizontal transmission were both negligible compared to adaptation to the floristic environment, we found that ancestry was more important than geographical proximity. Examples of markedly different plant use by peoples with opportunity for horizontal transmission highlight the importance of illness aetiology. For example, Shepard (24) describes two Amazonian societies sharing only 18% of species despite living in the same environment, and attributes these differences to contrasting illness theories. Cox (25) described Polynesian theories of disease causation, highlighting local supernatural beliefs, shared taxonomies of illnesses and a role for breaking social bonds. In this context aetiology would not be a barrier to transmission of knowledge, so the limited evidence for horizontal transfer of plant knowledge appears congruent with limited opportunities for cross-cultural transmission (18).

Against the background of adaptation to new floristic environments that we evidence here, we nevertheless sought to identify ancestral knowledge. To do so we compiled therapeutic applications, plant parts used, and local names for 98 shared plants: those used in more than one cultural group (Supplementary Data 5, 6 and 7 respectively). The use of cognate names and congruent (reconstructed) plant uses, including therapeutic applications and plant parts, are considered to evidence continuity of use (9). The peoples of the Bismarck Archipelago “voyaging nursery” are ancestors of the Polynesian voyagers (26). Reconstructing their knowledge, at the Proto-Oceanic (POC) node of our linguistic phylogeny, allows us to identify knowledge that accompanied the dispersal of Austronesian peoples into the previously uninhabited regions of Remote Oceania such as Polynesia. Twenty-two of the 98 plants for which we performed ancestral reconstructions had a probability of medicinal use at the POC ancestral node >0.5 (Table 1). Of these, four had ancestral reconstructions for both therapeutic application and plant part used with a probability >0.5, and two also had a reconstructed protoform name at the POC node. The names used for *Laportea* and *Thespesia* had protoforms that reconstructed more shallowly than the POC node, at the Proto Fijic (PFJ) and Proto-Eastern-Oceanic (PEO) nodes respectively, even though they had reconstructions for use at the POC node. Cognates descended from the protoform \**qaoa* are applied to all oceanic species of *Ficus*. *Ficus* as a genus had reconstructed ancestral use for digestive ailments and use of bark, leaf and root reconstructed at the POC node. Leaves of *Premna*, with protoform \**walo-walo* at the POC node, have reconstructed ancestral use for digestive and respiratory complaints. Figure 2 shows ancestral reconstruction at the POC node of use of *Premna*.

**Table 1.**
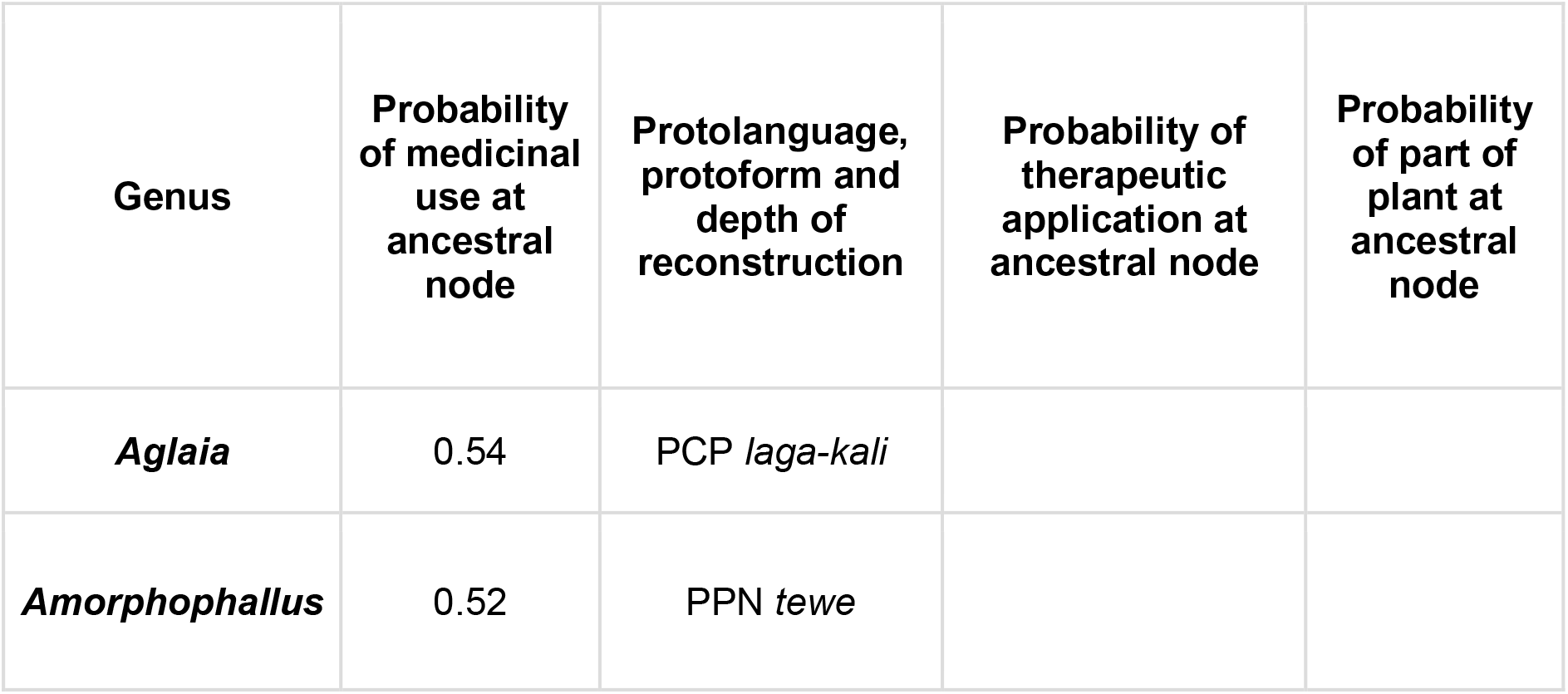
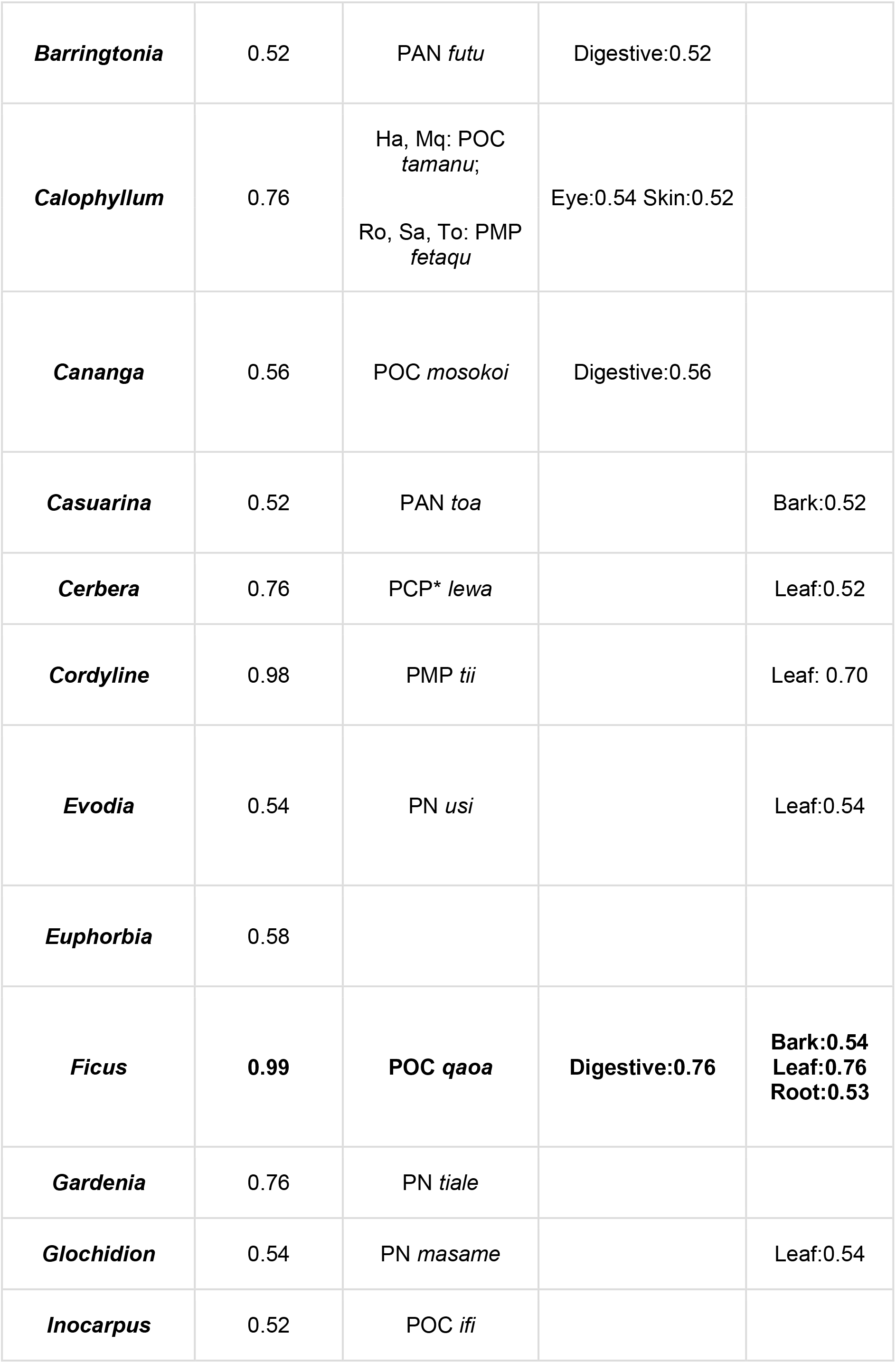
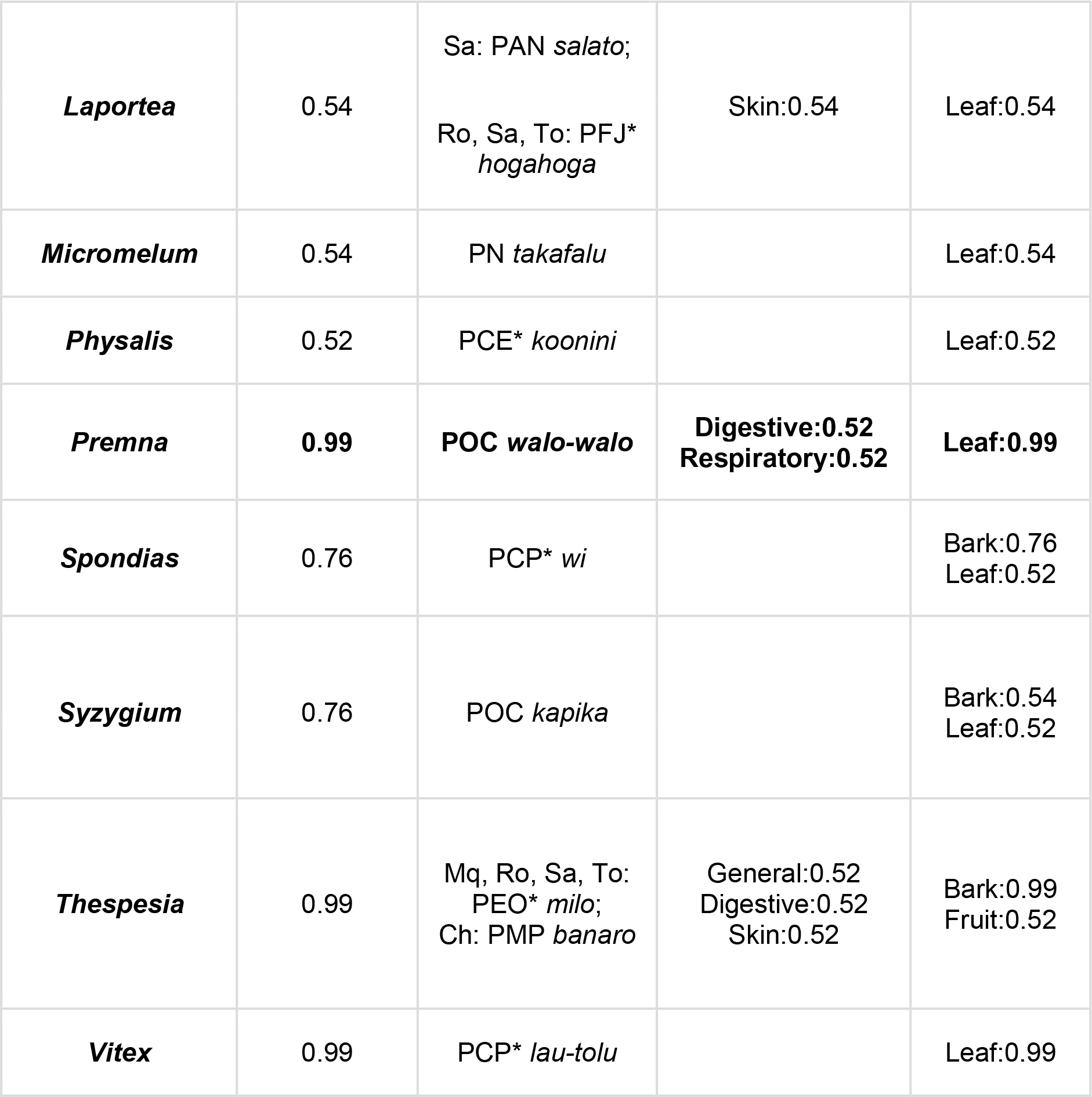
The list of plants at genus level for which the ancestral node for the six in-group cultures is reconstructed as having medicinal use with a probability >0.5. For these genera, protolanguages and protoforms are presented, and the diversity of therapeutic applications indicated. The protolanguage of the ingroup cultures is Proto-Oceanic (POC). Other protolanguages are as follows: PAN Proto-Austronesian, PMP Proto-Malayo-Polynesian, PEMP Proto-Eastern-Malayo-Polynesian (nodes deeper than the ancestral node for the in-group cultures); PCP, Proto-Central-Pacific, PEO Proto-Eastern-Oceanic, PFJ Proto Fijic, PPN Proto Polynesian (nodes shallower than the ancestral node for the six ingroup cultures, indicated with an asterisk). Protoforms are italicised. Therapeutic applications are according to the ICPC categorisation (A General, B Blood, blood forming, D Digestive, F Eye, H Ear, K Circulatory, L Musculoskeletal, N Neurological, P Psychological, R Respiratory, S Skin, T Metabolic, endocrine, nutrition, U Urinary, W Pregnancy, family planning, X Female genital, Y Male genital, Z Social), for each of the six cultures (Ch Chamorro, Ha Hawaiian, Mo Maori, Mq Marquesas, Ro Rotuman, Sa Samoan, To Tongan). The two plants with protoform, therapeutic application and plant part used reconstructed at or more deeply than Proto-Oceanic are in bold.

**Figure 2.**
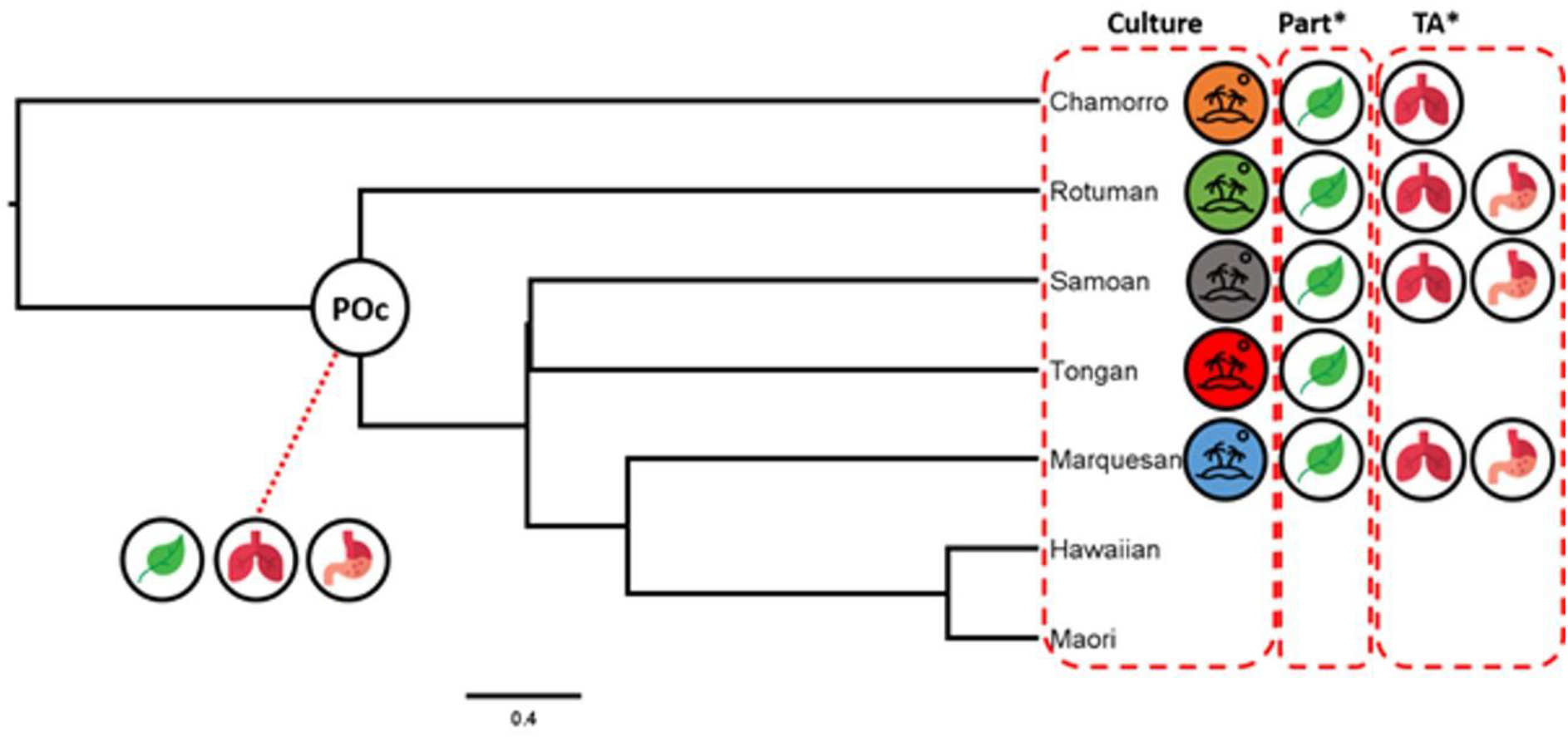
Reconstructed ancestral uses of *Premna*. Knowledge of leaves (*walo-walo*) for therapeutic uses for digestive and respiratory ailments is reconstructed to the Proto-Oceanic ethnolinguistic group.

Although we identified 22 plants that were used ancestrally, we found high levels of lability in the therapeutic applications and in the plant parts used. The degree of conflicting signal for plant part and for therapeutic applications are indicated by δ-scores and Q-residuals. The δ-scores and Q-residuals of 0.38 and of 0.01 for the therapeutic applications and 0.23 and 0.01 for plant parts used show that lability in therapeutic applications was particularly marked. Rectilinear webbing in the NeighborNet splits graphs also reveal lability in these aspects of use (Figure 3). Overall the degree of tree-like/web structure is comparable to other aspects of cultural knowledge, such as Eurasian folktale traditions (27), though less webbed than Austronesian music (28). Lability in therapeutic application could be attributed to the difficulties of scoring therapeutic applications for ethnomedicinal categories. Cox (25) noted the difficulties in relating Polynesian disease categories to Western categories, attributing this to the different views of disease causation. Others have also highlighted the difficulties in classifying therapeutic applications where there are incompatibilities between emic and etic views (29); this problem persists in phylogenetic studies and may be confounded by focus on systems of the human body rather than pharmacological properties (30). The Kruskal Wallis test showed that neither differing health needs nor cultural preferences for plant parts explained lability.

**Figure 3.**
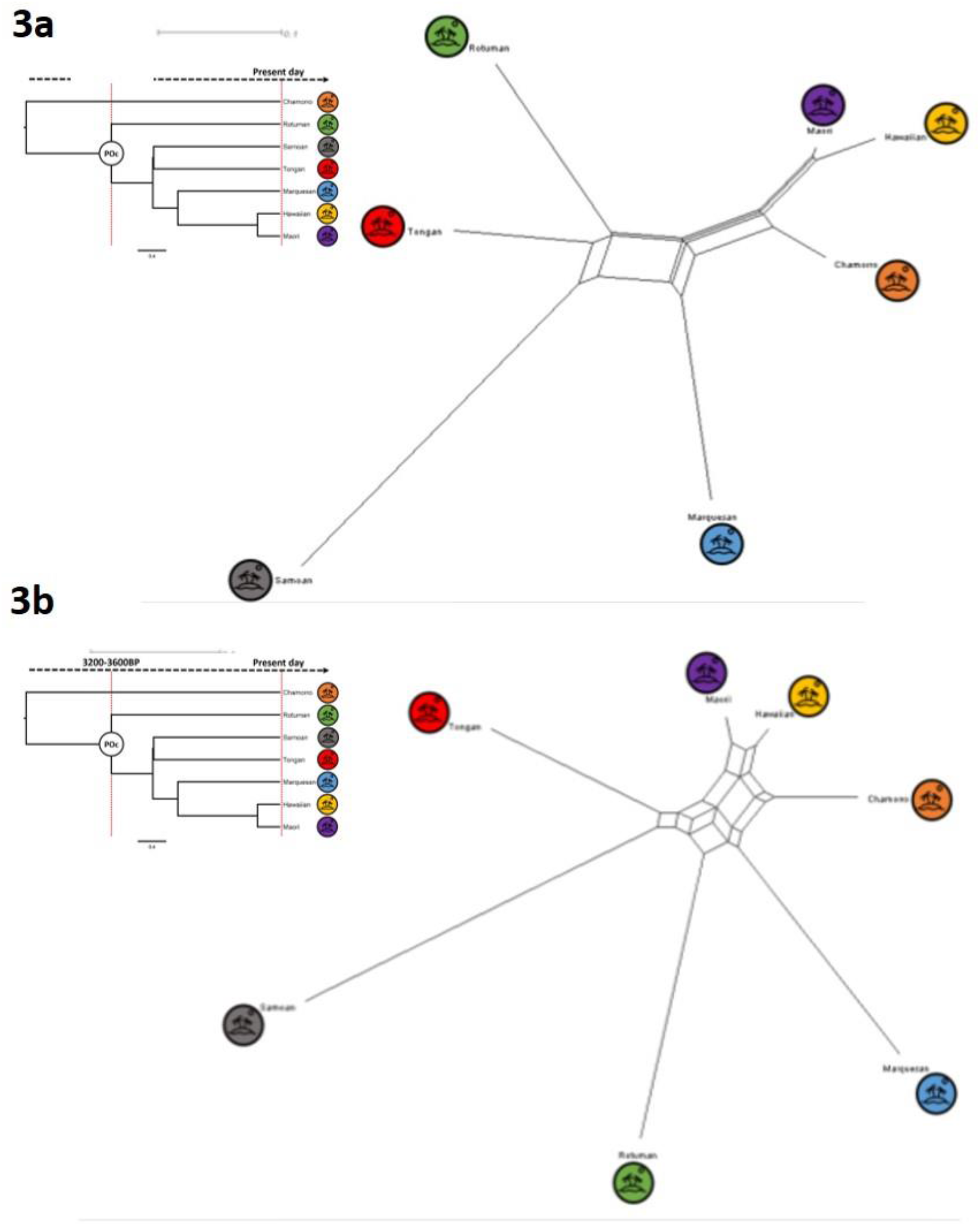
NeighborNet network visualizations of the conflicting signal in plant part used (3a), and therapeutic application (3b). Visualisations for the subset of 22 plants that are reconstructed as medicinal for the Proto-Oceanic ethnolinguistic group. Larger boxes indicate more conflicting signal. For comparison, linguistic relationships are shown as inserts in each figure.

We performed reconstructions for plants found in the local floras. We excluded plants thought to be recent, European introductions, though many such plants have been incorporated into Polynesian medicine (23). Whistler (31) tentatively classified 72 species as intentional introductions by Polynesians, introduced prior to European contact. These plants were identified because of their cultural importance and in many cases their distribution outside of Polynesia. Oral history recounts that these plants were introduced to Pacific islands by stowing seeds, cuttings, shoots, tubers and roots in canoes, and they are known as the canoe plants (31). Some of these species have well-established histories of dispersal based on phylogeographic study (32) whilst others remain the subject of ongoing controversy (33). Notably, *Premna* does not appear in the list of supposed canoe plants, and though *Ficus* does appear, it is considered by Whistler (31) to be introduced only to the atolls, and as native to our study sites. The putative native status of the plants suggests that migrants were carrying ancestral knowledge rather than the plants themselves. Our work might therefore be adding to the many examples of migrants encountering known species and retaining them in their ethnopharmacopoeias. An alternative scenario is that these plants were Polynesian introductions, but that medicinal plants are not sufficiently salient to participate in the “canoe plant” mythology.

Early missionaries to Polynesia did not describe established systems of plant medicine (23). This has been taken as evidence for a minimal historical ethnopharmacopoeia, or as a blindness to such systems amongst missionaries with little or no botanical interest but focussed on the healing practices involving the supernatural. Cox (25) suggests that if Polynesian medicine proved to be a pre-historic phenomenon and if this tradition appeared to be conservative, in the sense that the same plants were used for generations, this would be of much greater interest for Western medicine. Cox (25) also argues that Polynesian herbalism was an important component of early Polynesian cultures, perhaps descended from an earlier tradition brought by the first Polynesians to their new homelands. Our reconstructions point to relatively little continuity of use of ancestral medicinal plants. This might support the view that there was a minimal ancestral ethnopharmacopoeia. However, knowledge bottlenecks due to depopulation could also contribute to the low number of putative ancestrally-used plants. Terrible depopulation resulting from epidemics of influenza, smallpox, syphilis and others may have accounted for population declines of 98% in the Marquesas *(34, 35, 36),* 81% post-contact in Hawaii, and 60% in New Zealand (37). The limited retention of ancestral knowledge could be due to loss of opportunity for vertical transfer of knowledge in the context of depopulation. The potential effects of demographic change on cultural carrying capacity has been explored by cultural evolutionary scholars (38) and future work might model population decline with knowledge of plant uses.

Our study does not point to a significant role for translocation of medicinal plants, as is emphasised for plants used for food or material culture (31). Nor do we find a “relatively static interaction with the floristic environment”, considered by Schultes (39) to characterise knowledge of medicinal plants. Although a narrative over-emphasizing transformation may create the impression that living indigenous people are somehow “inauthentic,” having lost their connections to precontact roots (39), emphasising stasis creates a sentimental or mythic view of the holders of traditional knowledge (3). Our data contribute to the dismantling of the medicine myth, of traditional knowledge of plants tested by time and an integral part of culture passed on at deep timescales. Instead, we reveal both the creativity and adaptability of Polynesian people in meeting health needs on encountering new floristic environments.

## Materials and Methods

### Selection of study groups

A literature search was used to identify Austronesian ethnolinguistic groups with both an adequate documented ethnopharmacopoeia containing data of parts of plant, therapeutic applications and vernacular name, and a published checklist of angiosperm species present in the islands (generally archipelagos) that they inhabit. The data were compiled for Chamorro (Micronesian), Rotuman (Melanesian) and five Polynesian cultures, Hawaiian, Maori, Marquesan, Samoan and Tongan. The Polynesian selection reflects geographically widely dispersed cultural groups (Fig. 1a). The number of genera represented in each archipelago for which there was sequence data for phylogeny reconstruction ranged from 178 (81%; Tonga) to 747 (81%; Hawaii). The generic level phylogeny comprising 1,276 genera is presented in Newick format in Supplementary Data (4).

### Data preparation and scoring

Checklists of angiosperm species present in the islands or island archipelagos were sourced from published literature (Supplementary Data 1). Sources of the data describing medicinal plants use by the seven cultures are presented in Supplementary Data (2). Taxonomic names were checked and synonymy resolved using The Plant List (40) as a fixed source of accepted names. We compiled a matrix of the species used in at least two cultures that were not recent or European introductions according to Whistler (31, 41). This matrix included 98 shared species scored for absence (0) or presence (1) of medicinal use by each of the seven cultures. The absence (0) or presence (1) of use of parts (leaf, fruit, stem, sap, seed, whole plant, bud, bulb, fruit juice, pollen, bark, flower, shoot, petiole, rhizome, wood, gall) were scored from the original publications. Therapeutic applications were classified according to a modified International Classification of Primary Care (ICPC) standard (29). The absence (0) or presence (1) of therapeutic applications (General, Blood and blood forming, Digestive, Eye, Ear, Circulatory, Musculoskeletal, Neurological Urinary, Psychological, Respiratory, Skin, Metabolic including endocrine and nutrition, Pregnancy and family planning, Female genital, Male genital and Social) were also recorded. Vernacular names were sourced for all the shared species using the sources listed in Supplementary Data (2) and the comparative dictionary of Polynesian languages, POLLEX-Online (42). Coding of cognates, defined here as homologous words referring to the same plant species, was informed by multiple sources including POLLEX-Online. The presence or absence of words from each cognate set was coded as ‘1’ or ‘0’, respectively, to produce a binary matrix for 7 languages.

### Preparation of similarity matrices

Relatedness of cultural groups. A language phylogeny was a proxy for ancestral relatedness. The relatedness of the seven groups under study were derived from the time-calibrated MCC Bayesian phylogenetic tree for Austronesian languages sourced from D-PLACE (43) and inferred by Gray et al. (17). The tree was pruned to include the primary languages spoken by the seven cultural groups and exported (Figure 1b). A matrix of pairwise cultural distances was compiled using branch lengths to estimate evolutionary distances.

Relatedness of ethnopharmacopoeias and of floras. Community phylogenetic methods applied to a phylogeny of the flowering plants were used to calculate phylogenetic distances, first as an estimate of the relatedness of the ethnopharmacopoeias, and secondly the relatedness of the floras. Using the lists of genera, similarities between every pair of cultural groups were calculated for the plants in the floristic environment, and also for the medicinal plants. In order to calculate phylogenetic similarities, a phylogeny was reconstructed to include the angiosperm genera present in floras of all seven cultural groups. The phylogeny was reconstructed using nucleotide sequences for the photosynthetic enzyme ribulose-1,5-bisphosphate carboxylase oxygenase (rbcL), retrieved from Genbank for an exemplar specimen of every angiosperm genus present in the summed checklists. Sequences were downloaded using Geneious ver. 8 software (44). Wherever possible, the sequence downloaded represented a species present in the region, otherwise an exemplar from the same genus was selected. Sixteen non-Angiosperm species included as outgroups. Species names and accession numbers are presented in Supplementary Data (3). Alignment of rbcL sequences was implemented in Geneious, using Clustal W (45) and adjustments were made manually in Geneious. Phylogenetic trees were reconstructed through the CIPRES Science Gateway V.3.3 (46) using RAxML V8.2.10 (47) under maximum likelihood criteria. Subsequently, outgroups and problematic sequences (long branches, misplaced taxa) were pruned using the command “drop.tip” in ‘ape’ v5.3 package, R (48). The mean nearest taxon distance (MNTD) was used to assess floristic and medicinal similarities amongst the seven floras under null model 2, which randomly draws the sample taxa from the phylogeny pool. Phylocom (49) was used to make the calculations using the command “comdistnt”, with results presented in Supplementary Data (8).

Geographic proximity. We calculated geographical distance between all pairs of cultural groups using the Haversine formula, where the formula calculates the great-circle distance between two points on the globe, i.e. the shortest distance between them on the surface of the globe. This is appropriate for oceanic voyaging.

### Multiple regression

We used the four matrices to assess the contribution of proximity, ancestry and floristic environment to relatedness of medicinal ethnofloras. Multiple regression between medicinal plant selection as the response matrix and the three explanatory matrices, as follows:

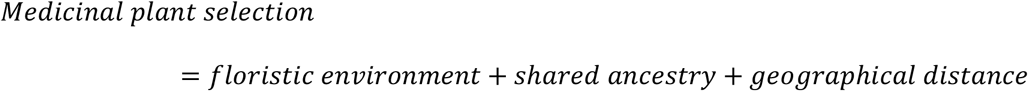

The MRM function in the R package Ecodist was used to perform the calculations, with 10,000 permutation tests to assess probability (50).

### Ancestral use

We consider ancestral use to be characterised by cognate names and congruent plant uses (plant part and therapeutic application) at an ancestral node (9). Firstly, we reconstructed absence (0) or presence (1) of medicinal use for each of the 98 shared species at the Proto-Oceanic node (POC; Figure 1a and 2). Ancestral states were reconstructed using Mesquite 3.01 (51) and maximum likelihood criteria, based on the Markov *k*-state 1 parameter model assuming equal rates of changes for both gains and losses. Reconstructions meeting a likelihood threshold of 50% were considered indicative of ancestral use. For the subset of 22 plants reconstructed as used, we reconstructed absence (0) or presence (1) of use of each plant part, and absence (0) or presence (1) of use for each therapeutic application using the same methods. We used POLLEX-Online (42) as a source of reconstructed protoforms and depth of reconstruction (protolanguage) for the vernacular names used by the ingroup cultures as recorded in the ethnomedicinal sources.

### Changing ancestral knowledge

We sought to visualise and quantify change in ancestral knowledge of medicinal plants, specifically variation in the use of the 22 medicinal plants belonging to the ancestral ethnopharmacopoeia. We used NeighborNet implemented in SplitsTree v4.14.6 (52) to visualize the strength of branching versus reticulate-like signals amongst the seven cultural groups for therapeutic applications and for plant parts used. We calculated two metrics for the therapeutic applications and plant parts data, δ-scores and Q-residuals (20) to estimate the degree of conflicting signal in these two aspects of plant use. These metrics were interpreted in the light of cultural preferences for plant parts and different health needs, estimated by calculating the number of reports per ethnolinguistic group of each plant part or therapeutic application. A Kruskal Wallis test was used to determine whether preferences or applications between cultural groups were significantly different.

## Supplementary data details

**Supplementary data 1.** Sources of flora checklists.

**Supplementary data 2.** Sources of ethnomedicinal lists.

**Supplementary data 3.** Species names and Genbank accessions.

**Supplementary data 4.** Newick-format phylogeny.

**Supplementary data 5.** Ethnobotanical linguistic data.

**Supplementary data 6.** Therapeutic applications of plants with shared use.

**Supplementary data 7.** Plant parts used of plants with shared use.

**Supplementary data 8.** Results of Phylocom analyses.

