## Supplementary data 1,2,4 and 8 for "Phylogenetic reconstructions of Polynesian medicinal plant use reveal adaptive strategies to meet health needs"

**Supplementary data presented here:**

**Supplementary data 1.** Sources of flora checklists.

**Supplementary data 2.** Sources of ethnomedicinal lists.

**Supplementary data 4.** Newick-format phylogeny.

**Supplementary data 8.** Results of Phylocom analyses.

**Supplementary data submitted separately:**

**Supplementary data 3.** Species names and Genbank accessions.

**Supplementary data 5.** Ethnobotanical linguistic data.

**Supplementary data 6.** Therapeutic applications of plants with shared use.

**Supplementary data 7.** Plant parts used of plants with shared use.

**Supplementary data 1. Sources of flora checklists**

| Flora | Source |
|---|---|
| **Hawaii** | Wagner, W. L., Herbst, D. R., and Lorence, D. H. Flora of the Hawaiian Islands website (2005). http://botany.si.edu/pacificislandbiodiversity/hawaiianflora/index.htm [Dec 2017]. |
| **Marquesas** | Wagner, W. L. and Lorence, D.H. Flora of the Marquesas Islands website (2002). http://botany.si.edu/pacificislandbiodiversity/marquesasflora/index.htm [Dec 2017]. |
| **New Zealand** | Saslis-Lagoudakis, C.H. et al. Phylogenies reveal predictive power of traditional medicine in bioprospecting. Proceedings of the National Academy of Sciences. 109, 15835-15840 (2012). |
| **Northern Mariana Islands** | Wagner, W. L., Herbst, D.R., Tornabene, M.W., Weitzman, A., and Lorence, D.H. Flora of Micronesia website (2012). http://botany.si.edu/pacificislandbiodiversity/micronesia/index.htm [Dec 2017]. |
| **Rotuma** | McClatchey, W., Thaman, R. and Vodonaivalu, S. A preliminary checklist of the flora of Rotuma with Rotuman names. Pacific science. 54, 345-363 (2000). |
| **Samoa** | Christophersen, E. Flowering plants of Samoa. (Kraus Reprint Co.,1971). |
| **Tonga** | Hemsley, W.B. The Flora of the Tonga or Friendly Islands, with Descriptions of and Notes on some New or Remarkable Plants, partly from the Solomon Islands. Journal of the Linnean Society of London, Botany. 30, 158-217 (1894). |

**Supplementary data 2. Sources of ethnomedicinal lists**

| Ethnolinguistic group | Source |
|---|---|
| **Chamorro** | Nandwani, D., Calvo, J.A., Tenorio, J., Calvo, F. and Manglona, L. Medicinal plants and traditional knowledge in the Northern Mariana Islands. Journal of Applied Biosciences. 8, 323-330 (2008). |
| **Hawaiian** | Nagata, K.M. Hawaiian medicinal plants. Economic Botany. 25, 245-254 (1971). |
| **Maori** | Saslis-Lagoudakis, C.H. et al. Phylogenies reveal predictive power of traditional medicine in bioprospecting. Proceedings of the National Academy of Sciences. 109, 15835-15840 (2012). |
| **Marquesan** | Girardi, C. et al. Herbal medicine in the Marquesas islands. Journal of ethnopharmacology. 161, 200-213 (2015). |
| **Rotuman** | McClatchey, W. The ethnopharmacopoeia of Rotuma. Journal of ethnopharmacology. 50, 147-156 (1996). |
| **Samoan** | Uhe, G. Medicinal plants of Samoa. Economic Botany. 28, 1-30 (1974). |
| **Tongan** | Whistler, W.A. Herbal medicine in the kingdom of Tonga. Journal of ethnopharmacology. 31, 339-372 (1991). |

**Supplementary data 4. Newick-format phylogeny**

(((((((Cycas:0.06824274416,((Nymphaea:0.05007665939,Trithuria:0.2175160684):0.04424692394,((((Hedycarya:0.02987405761,((((Cassytha:0.0816853801,((Beilschmiedia:0.0000010000005,Endiandra:0.006686754614):0.02251703682,Cryptocarya:0.03169332567):0.01112733734):0.0003314502326,(Cinnamomum:0.01079814377,((Litsea:0.006867526442,Persea:0.01861032834):0.01620665083,Laurus:0.005315709175):0.0052546162):0.02184999298):0.004915675167,Laurelia:0.03490749794):0.003797247759,(Gyrocarpus:0.04861629279,Hernandia:0.06667016917):0.01255253993):0.004159591814):0.01438098224,(Ascarina:0.06389504244,(Pseudowintera:0.06868960925,((Meiogyne:0.01054528158,(Cananga:0.0379562343,(Artabotrys:0.06277846407,(Annona:0.03902653907,Monodora:0.03099575495):0.01737164377):0.01588094572):0.00697280427):0.05335814529,((((Spirodela:0.006514185726,(Lemna:0.04361786368,(Landoltia:0.02628409881,Wolffia:0.08332262281):0.003540111029):0.02203509685):0.06256015298,((Anthurium:0.03518517382,(((Zantedeschia:0.1087783942,(Aglaonema:0.01302449784,Philodendron:0.02712352036):0.002633883097):0.001199942306,(((Xanthosoma:0.01755944895,(Syngonium:0.0129882695,Caladium:0.002325190905):0.004128946259):0.02183175529,(Amorphophallus:0.03772204651,(((Pistia:0.09188237683,Colocasia:0.0137129118):0.001780879484,Alocasia:0.01654229944):0.0000010000005,Arum:0.006535206797):0.01401997493):0.006388926434):0.01085662268,Cyrtosperma:0.02658162811):0.004635783053):0.003302558423,Dieffenbachia:0.05150527175):0.004301239178):0.004602519215,(Epipremnum:0.01048805192,Monstera:0.02217347342):0.01273032566):0.0192225586):0.04353726685,((Triglochin:0.0937631336,((Ruppia:0.07572602142,(Syringodium:0.04308313907,Halodule:0.0497138618):0.01036998725):0.02860797985,((Lepilaena:0.0396415176,((Potamogeton:0.02793575875,Stuckenia:0.02739806174):0.005711942546,Zannichellia:0.0427593668):0.001592812762):0.05282139193,Zostera:0.1104955975):0.04004504146):0.01333536671):0.03715963825,(((Najas:0.1445326313,((Vallisneria:0.07391101775,Hydrilla:0.06295472294):0.0302123307,(Halophila:0.05056778842,Enhalus:0.02189934886):0.1057661792):0.003198312841):0.01539887646,(Lagarosiphon:0.03974480063,((Egeria:0.009777935544,Elodea:0.02731816897):0.01371733731,Ottelia:0.0148930543):0.01095206641):0.0106944

8538):0.02005250592,Sagittaria:0.1579771759):0.02091551923):0.04697133262):0.01797451414,(((
Thismia:0.05129840487,(Dioscorea:0.06092216057,Tacca:0.05253227078):0.0204337591):0.036490
7532,((Ripogonum:0.07276371853,(Lilium:0.06271977084,Smilax:0.055356489):0.01105237852):0.0
212721186,((Gloriosa:0.02349718825,Iphigenia:0.02624803998):0.06891912721,Luzuriaga:0.03702
531806):0.05097875395):0.01191137482):0.0043466291,((Freycinetia:0.02431063103,Pandanus:0.0
1718457911):0.05523476491,(((((Eriocaulon:0.1958660422,((((Sporadanthus:0.01385953537,(Empo
disma:0.01835469137,(Apodasmia:0.02626417282,Centrolepis:0.1198616237):0.01233889609):0.01
640536555):0.08106126594,(Joinvillea:0.02609871996,(Achnatherum:0.00446157392,(((((Schizostac
hyum:0.002373054812,Bambusa:0.003849553543):0.0182247781,(Pseudosasa:0.01097752057,Phyll
ostachys:0.00225113108):0.01950579138):0.009575684942,(((Zotovia:0.004471171099,Microlaena:
0.007149691361):0.01645466352,Ehrharta:0.01252175981):0.02779957168,((((((Dichanthelium:0.0
1854073184,Entolasia:0.003624848231):0.012830391,(((((Setaria:0.03110308016,((Anthephora:0.01
254376959,Digitaria:0.0000010000005):0.005321169058,Echinochloa:0.03200880345):0.000001000
0005):0.003443791346,((Axonopus:0.02563090265,Paspalum:0.01463254012):0.01652196324,(Gar
notia:0.02204902881,(((((((Hyparrhenia:0.007086502096,(Themeda:0.01236280883,(Bothriochloa:0.
005282909607,Cymbopogon:0.03136988061):0.0000010000005):0.0000010000005):0.0030322401
02,((Pogonatherum:0.01510701778,(Imperata:0.003558740652,(((Eulalia:0.0000010000005,Sacchar
um:0.003526502971):0.0000010000005,Sorghum:0.003549158057):0.001723752508,Miscanthus:0.
001793891658):0.0000010000005):0.001730180994):0.002755589413,((Heteropogon:0.003508671
462,(Schizachyrium:0.001776581561,Dichanthium:0.005268769236):0.0000010000005):0.00000100
00005,(Andropogon:0.00533867295,Ischaemum:0.003563228672):0.001720613682):0.0000010000
005):0.0000010000005):0.004519780348,((Mnesithea:0.001952032966,Hemarthria:0.00530895423
4):0.003367892123,Coix:0.01599912804):0.0000010000005):0.0000010000005,Microstegium:0.005
380263014):0.0000010000005,Chrysopogon:0.007061940589):0.003734797927,Zea:0.00514757793
7):0.009580429033,Arthraxon:0.01696151916):0.0108212045):0.008849227524):0.001983819675):
0.03910397302,((((Cenchrus:0.0129488038,Stenotaphrum:0.007195549222):0.001729318153,Spinif

ex:0.0137730792):0.0000010000005,Ixophorus:0.01456631808):0.003401827548,Dissochondrus:0.007299915309):0.01368489897):0.005767744488,Sacciolepis:0.0200840136):0.0000010000005,(Oplismenus:0.01354591294,Cyrtococcum:0.006608599535):0.01076311933):0.0000010000005):0.002015990283,(Panicum:0.01887354911,((((Urochloa:0.00306955753,Megathyrsus:0.001527187277):0.0000010000005,Eriochloa:0.005354147672):0.002979230286,Melinis:0.01984869431):0.01060707686,Thuarea:0.002312371532):0.01153619188):0.009779642546):0.02122377412,Centotheca:0.04200728012):0.0000010000005,Lophatherum:0.02373576433):0.009399245078,((Arundo:0.03349117007,(Isachne:0.0261669306,Phragmites:0.01185883553):0.006134758477):0.004904755062,(((((Sporobolus:0.005988315049,Spartina:0.01469922872):0.005377853399,Zoysia:0.007830346482):0.009289712552,((Dactyloctenium:0.01230198835,(Tragus:0.01036363226,(((Chloris:0.01648294688,(Cynodon:0.007279585627,Lepturus:0.009817276656):0.001526563487):0.004894830584,Eleusine:0.01637958697):0.003424275956,Diplachne:0.01447154034):0.001548430536):0.001689712806):0.001660359825,((((Muhlenbergia:0.01874259032,(Disakisperma:0.01000309777,(Leptochloa:0.001642800684,Eustachys:0.008364113736):0.0000010000005):0.006800151382):0.001439946028,Distichlis:0.006631668964):0.001661829152,Bouteloua:0.02374790656):0.0000010000005,Eragrostis:0.001646996629):0.01556662736):0.001962712077):0.02609496177,Aristida:0.05677328031):0.01042223264,((Chionochloa:0.007644819883,(Cortaderia:0.003945355316,Pyrrhanthera:0.04205412552):0.02629374181):0.01782167946,Rytidosperma:0.02353442127):0.01707884726):0.01202830042):0.00269492179):0.03025174325,(Zizania:0.02323993662,Oryza:0.01949769792):0.01382658029):0.00595599169):0.003987291168):0.009024457679,(((((Triticum:0.007120452975,((Elymus:0.0000010000005,Bromus:0.02480986271):0.0252591303,Thinopyrum:0.0000010000005):0.01425765386):0.0000010000005,((Stenostachys:0.003960706545,Hordeum:0.008160356757):0.01439304317,(Leymus:0.007779134412,Australopyrum:0.01105824308):0.001743148528):0.007024990866):0.01210947352,Glyceria:0.03134303621):0.007858503718,(((((Poa:0.006559855875,Simplicia:0.007302760097):0.008935781461,(Alopecurus:0.01107387235,Phleum:0.01293441797):0.001736105892):0.003745914976,Puccinellia:0.03206183006):0.003197981766,(Arrhenatherum:0.01080612941,(((Sphenopholis:0.0074

51874836,(Trisetum:0.01887878886,Koeleria:0.0000010000005):0.008973539234):0.01335975608,
Phalaris:0.005318140307):0.02283439718,(Anthoxanthum:0.003632944544,((Aira:0.00368818243,((
Dichelachne:0.0000010000005,Echinopogon:0.0000010000005):0.0151800261,Calamagrostis:0.000
0010000005):0.003645153846):0.001810674648,(((Gastridium:0.009171448663,(Polypogon:0.0053
52960739,Lachnagrostis:0.001781359228):0.0000010000005):0.003489846554,Agrostis:0.00499752
5278):0.004624760132,Ammophila:0.001841625484):0.004550514577):0.004616641553):0.004611
464399):0.004711328102):0.01648417443):0.0000010000005,((Dactylis:0.003393356545,Lamarckia:
0.0000010000005):0.03844500859,(((Festuca:0.0000010000005,Lolium:0.005409055004):0.018232
21368,Holcus:0.01312146083):0.00300867509,(Deschampsia:0.01883105741,(Briza:0.01140685273,
Avena:0.008418595687):0.006371285851):0.001535434039):0.01380850196):0.004029794105):0.0
1548274471):0.01885083631,(Nardus:0.05015186059,(Nassella:0.04291551815,Austrostipa:0.01275
391639):0.009032134539):0.009277504933):0.02639863484):0.02823055572,Anemanthele:0.03890
668478):0.02095726536):0.0755268175):0.06819437267):0.02232927958,Flagellaria:0.0754701396
5):0.009309881142,(Xyris:0.3058049696,(((Cladium:0.01875548009,(((Scleria:0.07640313314,((((Car
ex:0.01420791763,Uncinia:0.02264130569):0.007122414247,Scirpus:0.005943300596):0.007228409
016,((Schoenoplectiella:0.03977537556,(Fuirena:0.02309363478,Eleocharis:0.023253867):0.007999
876586):0.001770566426,((Fimbristylis:0.03573617636,Bulbostylis:0.03097347233):0.00921794993
1,((Bolboschoenus:0.02037285149,((Ficinia:0.008386494493,Isolepis:0.02564975734):0.0109583376
7,Cyperus:0.04346534818):0.004433847876):0.003150367425,Schoenoplectus:0.0304293752):0.00
3311418777):0.0000010000005):0.006722513075):0.01288908776,Rhynchospora:0.05851151743):
0.009602646412):0.008058589402,((Oreobolus:0.03903636921,((Baumea:0.0000010000005,Macha
erina:0.0000010000005):0.02382823185,Lepidosperma:0.01888530655):0.01811637223):0.0054363
7168,(((Tetraria:0.04870977191,Schoenus:0.00910723325):0.02909068801,Gahnia:0.04456233695):
0.003725798241,Morelotia:0.02904702478):0.00208432267):0.004134588565):0.002530501895,Ca
rpha:0.07213933969):0.00385076493):0.02903274722,(Mapania:0.03406368823,Scirpodendron:0.0
963770167):0.0307319514):0.0459772642,(Luzula:0.07930618478,(Juncus:0.07557849189,(Marsipp

ospermum:0.0000010000005,Rostkovia:0.0188389625):0.02972733321):0.01395989263):0.0246979
0994):0.04256803915):0.01477058917):0.01058487408):0.0000010000005,((Typha:0.03498672631,
Sparganium:0.07920286896):0.01619474243,(Ananas:0.0101622158,Tillandsia:0.02047954646):0.04
246231073):0.005111468948):0.02356211815,((((Aneilema:0.030072459,Commelina:0.0522121476
5):0.02707653707,Murdannia:0.06395216862):0.02547804205,(Palisota:0.06363844668,(Callisia:0.0
2335390825,Tradescantia:0.03601603443):0.03475605495):0.008527026583):0.05369661778,(Eich
hornia:0.02374283684,Monochoria:0.0284633081):0.08595760932):0.03247882053):0.0119501148
7,(((Metroxylon:0.02902553773,Latania:0.01170685128):0.004822562441,((((Ptychosperma:0.0114
4407966,(Cocos:0.01428963671,(Pelagodoxa:0.01289134805,Roystonea:0.02653273188):0.0029094
41482):0.0000010000005):0.004796276464,((Adonidia:0.003274745437,(Veitchia:0.0000010000005
,Balaka:0.004851713466):0.003189807836):0.001598151816,((Rhopalostylis:0.008517056908,Archo
ntophoenix:0.001611996193):0.001608294082,((Areca:0.002711853173,Clinostigma:0.0100521104
4):0.005450025048,Dypsis:0.006385541769):0.001687733889):0.0000010000005):0.003195854121)
:0.004810111938,Phoenix:0.01406104094):0.004795632802,((((Washingtonia:0.006449909392,Cary
ota:0.0524905265):0.005040450776,Licuala:0.01969934387):0.0000010000005,Livistona:0.0032442
8718):0.002054436893,(Pritchardia:0.05547270376,Trachycarpus:0.007162475001):0.00092958455
79):0.002003657019):0.001910188853):0.03263593067,(Musa:0.02088469349,(((Canna:0.07228624
318,Costus:0.07608403225):0.01057206612,(Donax:0.02947483774,(Goeppertia:0.02153011231,M
aranta:0.02181656129):0.02722190998):0.01124176393):0.007002120027,((Zingiber:0.1099223138,
(Curcuma:0.007780647288,(Alpinia:0.02405810344,Hedychium:0.01530295719):0.008114880947):0
.00523697392):0.03973199899,Heliconia:0.01155683603):0.003729000033):0.006505429952):0.043
45807648):0.006129205497):0.005727252628,(((Molineria:0.06927119665,(Collospermum:0.02243
226308,Astelia:0.01161184644):0.009013869934):0.01433227718,(((Platanthera:0.02011840059,(H
abenaria:0.01908008418,Herminium:0.009812621405):0.003903937282):0.01222377598,(((Pterosty
lis:0.03288267605,Spiranthes:0.04386933387):0.001143832895,(Vrydagzynea:0.07301946006,(Ano
ectochilus:0.0150910229,(Zeuxine:0.1000439016,(Goodyera:0.01989280062,Platylepis:0.004194733

33):0.00222516472):0.002336775389):0.01337998263):0.008227060748):0.003339805668,((Adenoc
hilus:0.02035734772,((((Aporostylis:0.0185047228,(((Thelymitra:0.009318681522,Calochilus:0.0096
25749697):0.0108388159,Chiloglottis:0.01478703679):0.001922908073,Pyrorchis:0.00920413086):0
.0000010000005):0.003492240824,(Caladenia:0.06075659959,(Orthoceras:0.01324539838,Cryptost
ylis:0.031766652):0.003061461477):0.002224009724):0.004083794526,Didymoplexis:0.0313939232
3):0.003893400178,(Microtis:0.0189342296,Genoplesium:0.02045929604):0.00553332211):0.00200
871366):0.0000010000005,(Acianthus:0.01222376538,Corybas:0.03155500619):0.00634976563):0.0
02466611152):0.009813132399):0.01113625401,((((((Thrixspermum:0.02559178776,Papilionanthe:
0.007146565764):0.02989215979,(Glomera:0.01085124765,Arundina:0.009058392494):0.00763838
3887):0.003516443416,(Spathoglottis:0.01284508324,(((Malaxis:0.01509609961,Liparis:0.01501776
37):0.01655017771,(Oberonia:0.005067215515,Calanthe:0.007967795564):0.007401104462):0.008
874806898,((Coelogyne:0.01154206112,(Earina:0.02519934945,(Epidendrum:0.0113516249,Polysta
chya:0.03446421238):0.006378668926):0.005825440197):0.00425801882,(Pseuderia:0.0193736693
9,(Bulbophyllum:0.02210957384,Dendrobium:0.01591073149):0.005345470108):0.02191828244):0.
01027060263):0.003529712777):0.0000010000005):0.001786613166,(Eulophia:0.04003887149,((Ap
pendicula:0.03840593563,(Phreatia:0.02881179893,Trichotosia:0.0215923525):0.00288512385):0.0
06893743179,Phaius:0.0221427975):0.001837310183):0.003592524212):0.003333278707,Nervilia:0
.0841999727):0.009346884221,(Gastrodia:0.006785276214,Vanilla:0.09427578637):0.00792921569
8):0.005184163972):0.0383945551):0.001112892228,(((((Aloe:0.02303663609,Bulbinella:0.0288442
2151):0.01168997254,Asphodelus:0.02671183013):0.03521583808,(Herpolirion:0.0121785513,(Dia
nella:0.02248818481,Phormium:0.01368089725):0.006559783232):0.01046810387):0.01060272354
,((((Hosta:0.009277841609,Yucca:0.01426376037):0.001388295353,(Polianthes:0.00154414036,Aga
ve:0.003644664673):0.001570440432):0.02764831625,(((Beaucarnea:0.01157033802,(Sansevieria:0
.01973331406,Dracaena:0.01740429043):0.01915583537):0.01981129131,(Arthropodium:0.059473
34091,Cordyline:0.01138171519):0.01196695966):0.005681679601,Asparagus:0.04052116036):0.00
4369950724):0.004209709806,(Allium:0.0587335086,(Crinum:0.02451345031,((Eucharis:0.0059907

57141,(Pancratium:0.008779930948,Hymenocallis:0.006872586601):0.00690209127):0.0000010000
005,((Hippeastrum:0.00854814895,Zephyranthes:0.007281122066):0.0000010000005,Amaryllis:0.0
000010000005):0.01066614347):0.0132908526):0.005993455551):0.00714618486):0.00816185968
1):0.009422113861,(((Trimezia:0.02706255542,((Libertia:0.005086961818,Sisyrinchium:0.06366920
199):0.01814104698,Iris:0.02736944899):0.00569012625):0.01113132342,(Aristea:0.02781139485,(
Watsonia:0.04621350092,(Crocosmia:0.00730349445,Gladiolus:0.04243205344):0.007835053081):0
.01499059028):0.01640111906):0.02171800648,Xeronema:0.03423761067):0.001469986691):0.025
70275721):0.008704804757):0.007441645776):0.003041796947):0.02815160741):0.05414089097):
0.00514033253):0.007077232524):0.002758832366):0.001470126207,(((Peperomia:0.06195511025,
(Macropiper:0.01131758811,Piper:0.0150665341):0.02159070785):0.133055276,(Ceratophyllum:0.1
448724534,Aristolochia:0.1104389371):0.008788223571):0.01132071447,(((((Akebia:0.0611311740
7,(Berberis:0.06932238922,(Psychrophila:0.0380968233,((Anemone:0.01252472531,Clematis:0.023
54817105):0.03997088035,(Myosurus:0.0506441037,Ranunculus:0.02828414237):0.02030620154):
0.008978105595):0.03066468872):0.01137421611):0.002815616485,(Cocculus:0.01465665025,Step
hania:0.03791899888):0.0521930925):0.01020381445,(Fumaria:0.09006389292,(Argemone:0.04039
540872,(Hunnemannia:0.01322149568,Eschscholzia:0.01040053039):0.02611475322):0.004059240
761):0.009502171294):0.008167099647,((Banksia:0.02601891349,(Knightia:0.009385448473,(Maca
damia:0.05025002125,((Grevillea:0.03067411695,Hakea:0.01031016868):0.02889588235,Embothriu
m:0.0135395116):0.007600625401):0.01120670063):0.003921447335):0.00783229339,Persoonia:0.
04958070193):0.04541869352):0.004273172856,((((Ximenia:0.05937727128,(Anacolosa:0.0478367
6422,((Exocarpos:0.05673911503,Santalum:0.02308392775):0.01997470401,(Korthalsella:0.109006
635,(Loranthus:0.04003301609,(Peraxilla:0.02270878913,Tupeia:0.07145454299):0.001053434815):
0.07323879877):0.008658280351):0.01650993366):0.004916626563):0.01408699843,((Tetrastigma:
0.03636567551,(Vitis:0.007002743228,Cissus:0.02034412155):0.01172579637):0.06157480316,((((((
(Lumnitzera:0.02136419979,((Quisqualis:0.007012402164,Combretum:0.004402398847):0.0112808
1099,(Conocarpus:0.006124861924,Terminalia:0.04703010512):0.001943034425):0.007531167423)

:0.03872946975,((Ludwigia:0.03742037291,((Oenothera:0.02333546951,Epilobium:0.0375891106):0
.03172728529,Fuchsia:0.009338349389):0.02974747564):0.01688160036,(Lagerstroemia:0.0287497
5285,(((Ammannia:0.03735905758,Lythrum:0.05873145688):0.01021850378,Cuphea:0.0447036413
7):0.006979858226,(Pemphis:0.01500977661,Punica:0.02087033599):0.008628831821):0.00890769
3562):0.006446514319):0.0151203046):0.008338090039,((((Medinilla:0.03042882655,(Tetrazygia:0.
0232067927,Clidemia:0.01911525843):0.01403964213):0.001473027551,(((Tibouchina:0.011990135
24,(Melastoma:0.0133654824,(Heterotis:0.02546869297,Arthrostemma:0.02537848044):0.0027590
92481):0.001798980722):0.003185273708,Heterocentron:0.03704285918):0.008383296257,Pterole
pis:0.01069369078):0.006016431139):0.04270442122,Memecylon:0.05426271438):0.08107053042,
((((((Eucalyptus:0.008959368863,Angophora:0.01101653543):0.002234845374,(Kunzea:0.00902068
1281,Leptospermum:0.01449788777):0.01330133899):0.001777489324,(Metrosideros:0.016882679
95,Lophostemon:0.03017731375):0.002735312424):0.001691546536,(Melaleuca:0.01316853541,Ca
llistemon:0.004507586528):0.01563744554):0.001857329803,((((Decaspermum:0.01035798058,Ne
omyrtus:0.00436531536):0.006961963639,Lophomyrtus:0.006148821071):0.003448604173,Psidium
:0.01084289408):0.003095281871,Eugenia:0.01368367673):0.0117529926):0.002298587598,Syzygi
um:0.01746825623):0.01671071744):0.01198128382):0.03348190396,(((Muntingia:0.09182564624,
(Bixa:0.05865980017,(((Daphne:0.005463098562,Wikstroemia:0.02041258187):0.01991779096,(Ph
aleria:0.0202412329,(Drapetes:0.0412211975,Pimelea:0.02580867996):0.01590478405):0.0042626
49165):0.04740891746,(((Commersonia:0.01310153537,((Melochia:0.02850564198,((Triumfetta:0.0
1322214708,Heliocarpus:0.007081554839):0.03765917267,Waltheria:0.01920712272):0.025953274
61):0.01222833709,(Trichospermum:0.0389380001,Grewia:0.03003113077):0.01062562989):0.0026
79360821):0.0000010000005,((((((Pachira:0.0023988198,Ceiba:0.002166963991):0.008110870247,
Ochroma:0.005099506837):0.004719505144,(Heritiera:0.008690306784,Sterculia:0.01681125002):0
.001488871567):0.01002442622,(((Thespesia:0.01919650931,Gossypium:0.02467500015):0.010398
70305,(((Hibiscus:0.0106217681,Abelmoschus:0.03522343482):0.009159914773,Urena:0.00781449
8231):0.004787719649,(Abutilon:0.02009244322,Malvaviscus:0.01501713376):0.03730801356):0.0

2861412089):0.01088844244,((Malva:0.04889968228,Modiola:0.02017743879):0.011236719,((Hoh
eria:0.0000010000005,Plagianthus:0.003064870317):0.03112476353,(Sida:0.02219657512,Herissan
tia:0.01951511387):0.01942809586):0.0000010000005):0.008170428613):0.01007514198):0.01218
461137,Theobroma:0.009800237305):0.002132235796,Kleinhovia:0.01545071299):0.000001000000
5):0.006593895541,Entelea:0.03179591629):0.03982297245):0.007721439881):0.009056976131):0.
01147957226,((((Zanthoxylum:0.02674195933,((Ruta:0.05340551493,(Citrus:0.02703777237,(Tripha
sia:0.007727303697,(Murraya:0.03245761126,Micromelum:0.00764148799):0.004751544185):0.00
00010000005):0.03249557856):0.006585852016,((Euodia:0.05841622849,(Acronychia:0.015716108
8,Melicope:0.02614718417):0.006306539191):0.01880853283,Flindersia:0.03269115509):0.005390
192257):0.003478694902):0.02469610812,((Cedrela:0.005491696581,Swietenia:0.009229058938):0
.00125988208,((Melia:0.04474294601,(((Dysoxylum:0.02996608941,Aglaia:0.01417693881):0.01143
063743,Vavaea:0.0276494219):0.01413952382,Sandoricum:0.0204276075):0.02733501546):0.0150
4318244,(Toona:0.02411493852,(Xylocarpus:0.009986849346,Carapa:0.00194349858):0.007587429
021):0.00168896595):0.01645572788):0.0168314491):0.01731528438,((Pleiogynium:0.0042741200
59,((Dracontomelon:0.02760008941,Spondias:0.01644495447):0.004042045477,((Canarium:0.0215
2533953,Garuga:0.04758256631):0.03137497358,((Schinus:0.01394338965,Rhus:0.02761420778):0.
01103699014,(Mangifera:0.03016358996,Anacardium:0.05694413474):0.009564361785):0.0152542
2998):0.004974194956):0.004094569517):0.01877553217,(((Litchi:0.008124416251,(Pometia:0.016
1456468,Nephelium:0.005224948792):0.001449587382):0.0112124585,(((((Cupaniopsis:0.00843276
0809,Guioa:0.01817623724):0.002119894673,(Arytera:0.01430965603,Cupania:0.04048814484):0.0
04857687573):0.01175544666,Cardiospermum:0.06859617611):0.0000010000005,((Alectryon:0.01
186670475,Melicoccus:0.01695371135):0.006171593654,(Tristiropsis:0.01403971742,(Koelreuteria:
0.0437642279,Allophylus:0.03878185492):0.002150014737):0.01564831515):0.001817674227):0.00
2944917385,Sapindus:0.04318452173):0.0000010000005):0.01301423258,(Acer:0.04301522004,(Fil
icium:0.03028405733,(Dodonaea:0.0015816464,Harpullia:0.009774083142):0.01059962211):0.0044
63553891):0.003908111376):0.03243718118):0.001914920883):0.00911693366,Tribulus:0.0651410

8289):0.04166910839):0.00977086895,(Perrottetia:0.05431197395,(Tropaeolum:0.0447359747,((Batis:0.1492462069,((Gynandropsis:0.003402483786,((Cleomella:0.04372849553,(((Descurainia:0.01401946109,Pachycladon:0.02460320061):0.001937917726,(Capsella:0.01123030704,Lepidium:0.02437262871):0.001504354271):0.001961016388,((Rorippa:0.01629729362,(Nasturtium:0.01076947651, Cardamine:0.01013587374):0.004109968127):0.007172784106,(Notothlaspi:0.009341330851,(Lobularia:0.009058374102,(Sisymbrium:0.01308177315,(Hirschfeldia:0.01157235238,(Eruca:0.1269767648,(Rapistrum:0.04316969396,(Brassica:0.008239792244,Raphanus:0.009883988265):0.002852588458):0.008003606747):0.004879322544):0.009446269193):0.006142332659):0.003073799334):0.001489429175):0.002778522445):0.02123446609):0.009843175362,(Polanisia:0.01128738017,Cleome:0.01871902556):0.00568169895):0.004661094937):0.003460038685,Capparis:0.03647473074):0.04438692253):0.05686420042,(Carica:0.05406764076,Moringa:0.0210556386):0.01019089604):0.007188298674):0.01597334462):0.0006758989704):0.01737871902):0.00692686123,((((((Juglans:0.02145339838,(Casuarina:0.04781442706,(Betula:0.01391864757,Alnus:0.009925453891):0.00531755709):0.008529116762):0.005411275421,Quercus:0.04637212411):0.02009643788,Nothofagus:0.05788950427):0.01454191011,((((Ulmus:0.01374752012,Celtis:0.03232212372):0.05225969791,(((Parietaria:0.0426638737,(Boehmeria:0.01305603775,Pipturus:0.02699220316):0.01245973902):0.01176935269,((Pilea:0.04290176173,((Elatostema:0.02823315879,Procris:0.02440921041):0.01379418674,((Dendrocnide:0.03529865216,(Urtica:0.04623873844,Hesperocnide:0.03196927789):0.04386620756):0.002025340925,(Laportea:0.0303171116,Urera:0.04124687393):0.01861940329):0.01094642931):0.002826487255):0.01697694364,((Maoutia:0.01334180017,Leucosyke:0.04620357345):0.009742434055,Cecropia:0.01576509537):0.00167701683):0.001784681391):0.0189754519,((Gironniera:0.03274316835,((Morella:0.04561339465,(Trema:0.01858078084,Parasponia:0.0143679486):0.03729232827):0.004063705022,(Humulus:0.01259447583,Cannabis:0.01819017043):0.04719178706):0.007060987595):0.003029747168,(((Broussonetia:0.01512460287,Fatoua:0.04352161476):0.02017042236,(Ficus:0.02604078107,Castilla:0.03964547834):0.008527566516):0.01384986372,((Streblus:0.03697536358,Morus:0.008305246062):0.01222636919,Artocarpus:0.07745782174):0.01045977253):

0.03205490939):0.007388387229):0.02286659556):0.01105601842,((Frangula:0.0324505403,Rham
nus:0.0151891485):0.03664640557,(Elaeagnus:0.1062355544,((Pomaderris:0.02832987399,(Discari
a:0.01507497866,(Alphitonia:0.02370824213,Colubrina:0.02282679094):0.00478616946):0.0031451
15821):0.005078403743,Gouania:0.0549221103):0.02018727059):0.005339927543):0.00302916773
7):0.01121317701,((Rubus:0.02788593726,((((Fragaria:0.02788041788,Potentilla:0.06901685748):0.
0000010000005,Rosa:0.01564524294):0.005409795764,Acaena:0.05055236335):0.01249937725,Ge
um:0.09875459655):0.007045548936):0.04067034315,(Sorbaria:0.01443891375,(((Crataegus:0.001
709184176,((Photinia:0.0100282898,(Sorbus:0.008291301592,(Eriobotrya:0.001736376001,Cotonea
ster:0.008327033514):0.003436776654):0.0000010000005):0.001716930251,Pyracantha:0.0016825
03283):0.001663590459):0.001973076493,Heteromeles:0.02807861528):0.02852506253,(Spiraea:0.
03872207218,Prunus:0.05632328265):0.008371314827):0.004680840287):0.01124018458):0.02986
681529):0.01977443808):0.006458922407,(((((Ceratonia:0.01701023097,(Caesalpinia:0.0089083341
05,((Senna:0.01688003453,Chamaecrista:0.03360363346):0.009923377161,((((Adenanthera:0.0172
4581553,((Desmanthus:0.03011895402,Leucaena:0.01331489957):0.01214160459,((Mimosa:0.0300
3062115,Vachellia:0.02116607877):0.002574636381,((Pithecellobium:0.007324321254,Inga:0.0113
5692705):0.0000010000005,((Paraserianthes:0.003469162952,Samanea:0.003469897548):0.000001
0000005,(Albizia:0.00947759346,Calliandra:0.03028950528):0.007013243298):0.001745206824):0.0
06906128871):0.001756420262):0.02089488038):0.008461176445,(Parkinsonia:0.01380226286,Del
onix:0.01448852776):0.003904535146):0.002229080792,Cassia:0.0215641611):0.002229619045,(H
aematoxylum:0.02497400102,Guilandina:0.01546342767):0.01838649012):0.002078587019):0.001
914071139):0.00439678627):0.00864556749,((Sophora:0.02688750033,((((Abrus:0.08039306271,(
Wisteria:0.04708755553,((Montigena:0.01219661683,(Carmichaelia:0.004638073471,Clianthus:0.00
3617639912):0.008505189065):0.06260396712,((Lathyrus:0.01094351869,Vicia:0.02303182587):0.0
4136561724,(Melilotus:0.03539004892,(Trifolium:0.02873299006,Medicago:0.02676990541):0.025
19228835):0.01140385205):0.03788752626):0.009496499194):0.01310016518):0.01321757167,(Lot
us:0.09983403393,(Sesbania:0.05049983518,Robinia:0.05346947186):0.00468127384):0.002739116

68):0.01093927652,((((((Macrotyloma:0.031986614,(Vigna:0.03339335181,((Dolichos:0.0020665817
26,Dipogon:0.001828322662):0.006120104721,((Phaseolus:0.004584265975,Sinapis:0.00574111078
2):0.02383518341,(Macroptilium:0.01306195866,Lablab:0.009466377389):0.003822469793):0.0015
93763112):0.001756885222):0.005027642302):0.02076129281,((((Neonotonia:0.02232692887,(Glyc
ine:0.0411589619,Pueraria:0.006470863888):0.004326580616):0.00565982379,Neustanthus:0.0152
3998953):0.001958474529,(Psoralea:0.02172074345,(Pachyrhizus:0.00551798451,Calopogonium:0.
01794573161):0.004653857432):0.008522476655):0.004997809428,Strongylodon:0.04863050961):
0.0000010000005):0.0192508809,(Erythrina:0.06302367458,(Flemingia:0.009141649537,(Rhynchosi
a:0.02719386129,Cajanus:0.01707465416):0.01032918222):0.02880767359):0.01086175187):0.018
57355887,((Mucuna:0.03444409199,((Dendrolobium:0.04405417264,(Uraria:0.0000010000005,(Aly
sicarpus:0.03422478818,Desmodium:0.02999290699):0.007721563963):0.00964244142):0.0124760
9769,(Lespedeza:0.00765072159,Kummerowia:0.0294059732):0.03827729258):0.0255808074):0.01
097378901,Kennedia:0.02652593452):0.00453830084):0.02794576525,((((Tephrosia:0.0393917122
2,Millettia:0.002702320618):0.004539625594,Derris:0.01128690796):0.01631341668,(Galactia:0.01
752492244,(Canavalia:0.01383893113,Dioclea:0.008942226401):0.01500432228):0.03734882103):0
.03453822258,(Clitoria:0.08765288265,Centrosema:0.04262584652):0.0137756362):0.00307825347
7):0.01224950699,(Entada:0.006559985074,Indigofera:0.009034241054):0.05951694635):0.005648
902053):0.007278330369,(Dalea:0.06696464481,(Zornia:0.0369804552,((Aeschynomene:0.0554501
1665,(Stylosanthes:0.0496016464,Arachis:0.01920192551):0.04418185127):0.01907514712,Inocarp
us:0.02424339561):0.009583289899):0.009756031619):0.003852294861):0.0000010000005):0.003
338705593,(Crotalaria:0.06939677517,(Podalyria:0.02685507195,(Cytisus:0.005018730814,(Ulex:0.
0199098404,(Lupinus:0.0159640021,(Genista:0.01248739643,Spartium:0.004421272263):0.002122
130086):0.0000010000005):0.004696023647):0.04977146076):0.0071907814):0.01896183898):0.01
315040831):0.006718685312,((Bauhinia:0.0206237145,Barklya:0.04483049652):0.02707675443,((C
ynometra:0.009293509273,(Saraca:0.0102624227,Intsia:0.004335174487):0.0000010000005):0.002
715342283,Tamarindus:0.0109394279):0.04537785316):0.001813475973):0.0105049065,(Suriana:0

.05967928465,Polygala:0.09675235023):0.004496457048):0.02639822253,(Coriaria:0.04036342255,((Begonia:0.04848366887,Hillebrandia:0.01788151587):0.02576820389,(Momordica:0.02636174352,(Bryonia:0.05491160809,(((Benincasa:0.01199466664,(Citrullus:0.003028316983,(((Cucumis:0.02886070484,Lagenaria:0.01309738559):0.002518956339,(Coccinia:0.006520464399,Melothria:0.03263330413):0.001778085259):0.002088804418,Zehneria:0.02074939123):0.001864093746):0.001377288844):0.009135894476,Cucurbita:0.01362265032):0.004679331881,(Luffa:0.00394147068,(Sechium:0.005109539345,Sicyos:0.01342855226):0.02112068391):0.00662959171):0.0000010000005):0.0110935662):0.04827923523):0.008232834005):0.02462166969):0.005861587027):0.003601969991,((((Putterlickia:0.006009967281,Gymnosporia:0.01049850951):0.007027992536,(Brexia:0.02064113298,(Maytenus:0.01107190847,((Stackhousia:0.1822585129,Euonymus:0.01205676266):0.006291325403,Celastrus:0.02747919785):0.005308787956):0.0000010000005):0.004320241895):0.06881618948,((Averrhoa:0.01461144363,Oxalis:0.1447852551):0.01509266866,(Spiraeanthemum:0.02982816171,((Weinmannia:0.02980803566,Caldcluvia:0.02612169158):0.01782295898,(Aristotelia:0.02437926494,Elaeocarpus:0.04664921353):0.008883093644):0.009807572809):0.007812378643):0.02108238542):0.001142122999,((Atuna:0.009218460174,(Chrysobalanus:0.008693606441,Parinari:0.01087734744):0.0000010000005):0.08877858095,(((Hypericum:0.1635499036,((Clusia:0.02182417566,Garcinia:0.05201618708):0.04078749602,(Calophyllum:0.008628489252,Mammea:0.0263406079):0.01761181521):0.00563295919):0.02852431278,(Linum:0.1234307327,(((Bischofia:0.02965829212,(Baccaurea:0.05057372732,Antidesma:0.03388045437):0.01290254593):0.01668538661,((Poranthera:0.08099981657,Flueggea:0.01967134455):0.02469360184,(Glochidion:0.01173185405,(Breynia:0.01200722376,Phyllanthus:0.01848349444):0.006991058384):0.05100104446):0.0168051487):0.01034938519,((Galphimia:0.03788055719,(Tristellateia:0.02910557818,((Hiptage:0.01422227695,Malpighia:0.01610922331):0.006055710904,Heteropterys:0.0190151548):0.004559294604):0.01095826724):0.03671716177,(Ochna:0.0424958529,Sauvagesia:0.06247470584):0.03383974453):0.003632622877):0.002521361113):0.002998764292):0.003934795885,((Elatine:0.09716931239,Drypetes:0.1352848635):0.007899701497,((((Pangium:0.06536441807,(Erythrospermum:0.04067969481,((Passiflor a:0.1016449234,Turnera:0.1135674861):0.06037199657,(Viola:0.09010031434,(Melicytus:0.023944 03831,Isodendrion:0.05038854726):0.00916099677):0.04876927187):0.01088327687):0.004863698 043):0.0000010000005,(Casearia:0.06136964357,(((Flacourtia:0.01810092338,Xylosma:0.01048241 07):0.009357976528,(Dovyalis:0.04382820452,(Populus:0.009304046039,Salix:0.05264148068):0.02 869577):0.003476459027):0.002262089487,Homalium:0.03960512458):0.03357096283):0.0168696 7931):0.00212964783,(Endospermum:0.03761455081,((Cnidoscolus:0.01503721427,Manihot:0.006 562148891):0.03534938222,(((Codiaeum:0.02997841001,Aleurites:0.00561699046):0.01587005495, (Croton:0.07341584401,Jatropha:0.008994008127):0.0135120565):0.009377857091,(((Homalanthus :0.01896355536,Excoecaria:0.01415299368):0.009029185088,(Pedilanthus:0.03672684838,Euphorb ia:0.02108741381):0.03274772353):0.01110646371,((Ricinus:0.01599143612,Melanolepis:0.014220 72716):0.005023305634,((Macaranga:0.01850355887,Mallotus:0.01546401787):0.03524625438,(Ac alypha:0.08205197519,Claoxylon:0.05807759775):0.003667631135):0.003490650827):0.009048298 767):0.005400142766):0.005375133236):0.0003279841252):0.02216077149):0.003266572144,(Cros sostylis:0.02076073628,(Rhizophora:0.03505169832,Bruguiera:0.01953559192):0.006260192504):0. 04633206897):0.0000010000005):0.003985689053):0.001660032643):0.02068970223):0.00857947 7347):0.009084218474,Ixerba:0.04961275609):0.005181187125):0.002664640608,(Pelargonium:0.0 5504503434,(Geranium:0.03269563132,Erodium:0.0442288535):0.04392478316):0.07887452015):0 .006742978636,(Ribes:0.1019408543,((Myriophyllum:0.02662015601,(Gonocarpus:0.02370988401, Haloragis:0.0148929263):0.002196115669):0.0654379693,((Crassula:0.1069160118,Sedum:0.04373 263905):0.01239935315,(Bryophyllum:0.0000010000005,Kalanchoe:0.0000010000005):0.02782752 067):0.0717031483):0.01434465046):0.0172654254):0.001896496635):0.005168711593):0.0042054 48655,((Dillenia:0.08876760206,((Drosera:0.1345227117,(((Plumbago:0.1019025174,Rumex:0.0820 3683216):0.006931720902,(Muehlenbeckia:0.0464249747,Fallopia:0.04648273509):0.03599886826 ):0.003193582567,((Persicaria:0.07074713223,Coccoloba:0.03060651617):0.02149935631,Antigono n:0.04013271074):0.01034227184):0.09424644623):0.01699897521,((((Polycarpon:0.05166271454,( ((((Vaccaria:0.06970177533,(Dianthus:0.007391938346,Petrorhagia:0.04029748635):0.0081834025

03):0.005534241711,((Cerastium:0.04604651278,Stellaria:0.0282152656):0.04924493229,Silene:0.03879651268):0.001112732177):0.002914260765,(Spergula:0.027904061,Spergularia:0.05738537546):0.008706871601):0.001364407585,(Arenaria:0.01175028154,(Colobanthus:0.005639494615,Sagina:0.01188152767):0.02402401235):0.01301863061):0.003402884408,((Drymaria:0.0964032671,Scleranthus:0.02934832767):0.04210717175,Schiedea:0.03752300985):0.02022854697):0.01549447276):0.02805529843,(((Suaeda:0.0403437327,Salicornia:0.05300225159):0.008853906488,Bassia:0.03726019684):0.026348265,((Salsola:0.04133483252,((((Philoxerus:0.02150738501,Gomphrena:0.01819627038):0.01098894893,(Nototrichium:0.0000010000005,(Alternanthera:0.05194180561,Achyranthes:0.0000010000005):0.01201823608):0.01629255283):0.02889131506,(((Cyathula:0.02543898549,Deeringia:0.01835202533):0.04483276166,Amaranthus:0.05449164877):0.01890008762,Celosia:0.04181503472):0.01739479207):0.01094924869,Charpentiera:0.03178038937):0.02145630968):0.029223844,(Dysphania:0.05938318034,(Atriplex:0.01826908046,(Rhagodia:0.0000010000005,Chenopodium:0.01226536625):0.02877836499):0.01840336987):0.008907350631):0.004863231288):0.02407544421):0.009168757752,((((Hectorella:0.0129734422,Montia:0.03335190582):0.01539040847,(Portulaca:0.04116671937,(Talinum:0.03205099869,Opuntia:0.04836036193):0.02898708694):0.004099275218):0.001700173704,Anredera:0.03969924246):0.008645216324,((((((Mirabilis:0.01488253319,Boerhavia:0.05918010687):0.02934406038,(Pisonia:0.002844253569,Bougainvillea:0.02257084471):0.00623785728):0.007345236632,Rivina:0.04990020598):0.01380494364,(Phytolacca:0.06497270023,(Sesuvium:0.06744420825,Trianthema:0.04104045299):0.04860940776):0.01284022343):0.003234447418,(Tetragonia:0.02544652431,(Carpobrotus:0.0000010000005,Disphyma:0.00481755439):0.04763069147):0.009220014664):0.01141590913,Mollugo:0.0958339195):0.009549952751):0.04167092207):0.05838020914,(Frankenia:0.07265518171,Myricaria:0.05387194995):0.0438611782):0.007686514568):0.06075119956):0.006581373216,(((Cornus:0.06041707177,(Hydrangea:0.06497738363,Philadelphus:0.01655469685):0.01628819683):0.009110035303,(((Maesa:0.03418370992,(((Ardisia:0.04586253456,Elingamita:0.0135631087):0.009485318614,(Rapanea:0.02135833818,Myrsine:0.003519285711):0.01307942877):0.003130030325,Lysimachia:0.03155515262):0.04316026655)

:0.03069814215,((Eurya:0.03691896154,Camellia:0.0266112613):0.01972350652,((((Rhododendron:
0.005829196306,(Erica:0.06174754329,Calluna:0.08724104747):0.006830630953):0.02567733775,((
Archeria:0.01488963634,(Dracophyllum:0.01326212912,((Pentachondra:0.04647213548,((Cyathode
s:0.03579056401,Leptecophylla:0.01328522756):0.0000010000005,(Leucopogon:0.02473809079,St
yphelia:0.02333284851):0.01164141932):0.007199247136):0.01207567549,Epacris:0.03194843294)
:0.006545541955):0.001686165874):0.01852827242,(Vaccinium:0.04127189524,Gaultheria:0.03331
889617):0.0219215683):0.03133275138):0.04069219439,Actinidia:0.05661897177):0.01386324425,
((Sideroxylon:0.01476142067,(((Isonandra:0.04649263241,Chrysophyllum:0.01084467493):0.05217
195602,(Planchonella:0.003869419656,(Palaquium:0.05052241233,Manilkara:0.0000010000005):0.
002475621257):0.001239587506):0.001805267304,Mimusops:0.05210323626):0.004525740409):0.
01833031611,((Cobaea:0.07403474911,Diospyros:0.04029123323):0.005536092033,Barringtonia:0.
05040276851):0.002000199433):0.001557381394):0.0000010000005):0.001361526209):0.0098546
36766,Impatiens:0.09615849832):0.009735860626):0.0119151504,((((Gomphandra:0.1039570389,((
(((((Sebaea:0.06081263081,Centaurium:0.03700715484):0.01332303373,Fagraea:0.0479678853):0.0
1134625953,Gentianella:0.05440089857):0.03614300435,((Logania:0.006503431516,(Geniostoma:0
.02573088646,Mitrasacme:0.03660460086):0.005066197111):0.02724147685,(Alstonia:0.01299021
147,(((Melodinus:0.02041021054,((Pteralyxia:0.01623674901,Lepinia:0.02419022917):0.009672329
478,Alyxia:0.01344417064):0.0390832058):0.00108018606,((((((Araujia:0.01587889117,Tylophora:0.
0130920103):0.003096476315,(Asclepias:0.0100319786,Calotropis:0.00568292193):0.00433963782
5):0.005333692805,(Stapelia:0.03159960477,(Marsdenia:0.02595241535,Hoya:0.01165595572):0.0
06295513762):0.003671513941):0.02022218175,(Strophanthus:0.01086864985,(Cryptostegia:0.036
22372095,Parsonsia:0.01502116606):0.007542865999):0.006405187684):0.002045786643,(Adeniu
m:0.006482375059,Nerium:0.004808907434):0.0000010000005):0.01556179628,((Allamanda:0.016
06157588,Plumeria:0.02121591411):0.0187213262,(Thevetia:0.0180266459,Cerbera:0.0148099920
3):0.01376483167):0.0111685082):0.001876635218):0.02739707466,((Ochrosia:0.02027785969,(Ra
uvolfia:0.02067808443,(Catharanthus:0.01063814685,Vinca:0.05193541138):0.0154679529):0.0031

62003613):0.01773159358,Tabernaemontana:0.0380699824):0.01167407335):0.004866433885):0.0
2470525622):0.005220001039):0.01120553058,(((Mussaenda:0.03213608982,((Aidia:0.0139210832
2,((Randia:0.02633351263,Coffea:0.04423788753):0.004311310769,(Gardenia:0.02707984029,Tare
nna:0.03764669498):0.007826595641):0.01060694557):0.0207522998,(Ixora:0.0597105213,(Psydra
x:0.004087203238,(Canthium:0.02009514052,Vangueria:0.02049179471):0.007134872959):0.01170
335886):0.00883790941):0.01189865186):0.0217036345,((((Coprosma:0.02071490676,Nertera:0.01
387620582):0.0179983024,((Pentas:0.06343998319,(Hedyotis:0.05767934443,((Diodia:0.012721005
81,((Mitracarpus:0.01374890689,Richardia:0.01094873402):0.0000010000005,Spermacoce:0.01634
397358):0.007756407158):0.02318901598,Oldenlandia:0.03148462849):0.01971631691):0.0343126
1929):0.02295211709,((Sherardia:0.04097354838,Galium:0.04571336398):0.07008465595,Serissa:0.
03605850849):0.007173489512):0.01003732168):0.006207915346,Paederia:0.02196288157):0.023
03410366,((Psychotria:0.03782346213,Geophila:0.04705939725):0.02855175648,(Gynochthodes:0.
02577547924,Morinda:0.01731032276):0.008791950224):0.01106347015):0.05639553217):0.00775
8944086,(((Guettarda:0.006448305875,(Bobea:0.008120836239,Timonius:0.001801205441):0.0033
13428514):0.01109329533,((Bikkia:0.00122571756,Badusa:0.01775539743):0.02874163483,(Naucle
a:0.01064704337,Neonauclea:0.008498923748):0.02299677229):0.002103054163):0.003758954374
,Cinchona:0.02447372811):0.02167824841):0.02063253531):0.03040366931,((((((Nestegis:0.009725
370073,(Jasminum:0.05009802526,Noronhia:0.01336652951):0.004800800634):0.005366765593,Ol
ea:0.007926162216):0.001873739862,Chionanthus:0.0118868063):0.005457410238,(Fraxinus:0.011
81761808,Ligustrum:0.014203122):0.005363721086):0.02180974317,(Cyrtandra:0.118580499,(Rha
bdothamnus:0.02213326323,(Jovellana:0.03047754461,(((((Buchnera:0.0779174191,Angelonia:0.04
864800444):0.01530501018,(((((Premna:0.03724184328,((Teucrium:0.01411238587,(Ajuga:0.01952
767767,(Oxera:0.01507085121,Clerodendrum:0.02931837996):0.02126838579):0.006903425982):0.
01566479075,Tectona:0.01388240645):0.003835289747):0.005879575283,((Ocimum:0.0262786090
5,((Lepechinia:0.001765499024,Prunella:0.02433158679):0.001759223728,((((Plectranthus:0.00835
0340593,Leucas:0.0130810151):0.003357985117,(Condea:0.01227078446,(Cantinoa:0.0073242329

11,Hyptis:0.009211594356):0.02270595425):0.007317754057):0.01299014695,(Mentha:0.00684561
134,Thymus:0.01536933595):0.01559373737):0.006994885876,Salvia:0.03333691379):0.015339585
07):0.01396286355):0.03649495499,(((Pogostemon:0.06217899311,(Lamium:0.02631299896,(((Leo
notis:0.02301110612,Marrubium:0.01224503145):0.006713724827,Leonurus:0.01035126223):0.005
731042458,((Haplostachys:0.0000010000005,(Phyllostegia:0.003513683856,Stenogyne:0.00000100
00005):0.001749040691):0.01935504355,Stachys:0.02223478707):0.00663937146):0.00679239180
7):0.001948790766):0.007778081237,Scutellaria:0.03984353852):0.00539875972,Vitex:0.02623123
044):0.003726674824):0.009206822471):0.01491095494,(Torenia:0.03031955266,Lindernia:0.0465
4306664):0.0263510716):0.001302914606,(Myoporum:0.05036032225,((Thunbergia:0.0499140529
2,Avicennia:0.03878456474):0.005848645224,((Acanthus:0.02173694682,Crossandra:0.0193183518
2):0.03650855018,(Barleria:0.04220114099,((Asystasia:0.0399820754,((Pseuderanthemum:0.01178
859975,Graptophyllum:0.0000010000005):0.01165632853,(Justicia:0.01766822513,(Dicliptera:0.02
819194077,Hypoestes:0.01592892388):0.0136695686):0.0110087708):0.003250045387):0.0246297
1149,(Ruellia:0.01794241127,(Eranthemum:0.02684888823,(Sanchezia:0.01631916011,Hemigraphis
:0.02779147341):0.01950382242):0.0000010000005):0.0306972435):0.001802522317):0.01721482
992):0.0000010000005):0.01324488665):0.005824098947):0.002125776441,((Spathodea:0.0186681
1757,((Kigelia:0.01431359924,(Crescentia:0.01925704044,Dolichandra:0.03445519595):0.00153550
5858):0.003146100256,(Jacaranda:0.02179374907,(((Tecomaria:0.009819388347,((Tecomanthe:0.0
03354861284,Pandorea:0.003979436366):0.001824985328,Podranea:0.005437773177):0.00000100
00005):0.00365681866,Tecoma:0.007984198197):0.009313087122,Catalpa:0.007241662925):0.002
940504316):0.0000010000005):0.001461488544):0.001424717822,Tabebuia:0.005951053124):0.00
8501521243):0.008770337862):0.0000010000005,(Bacopa:0.0321357962,(Gratiola:0.05442557055,
Dopatrium:0.04859338012):0.01489600549):0.01543185051):0.0000010000005,((Utricularia:0.0881
1472501,(Tetrachondra:0.03453527936,Glossostigma:0.04146133024):0.00368742958):0.00249548
9085,(((((Linaria:0.03007702752,Lophospermum:0.03344884172):0.003947391534,Cymbalaria:0.00
6762334712):0.008518494662,((Veronica:0.05173091523,Plantago:0.08643542012):0.01809404289

,Callitriche:0.07409201259):0.007763839953):0.01683865,Sesamum:0.01870924533):0.0037609205
93,(Mazus:0.05144863478,((Buddleja:0.02032478461,Verbascum:0.01590267282):0.00863020364,((
Mimulus:0.004403617452,Citharexylum:0.0394122084):0.0000010000005,(Verbena:0.04906082374
,((Phyla:0.03530608573,Lantana:0.01647336453):0.03493261355,Stachytarpheta:0.03036680189):0
.01262811722):0.006297393065):0.01296017299):0.003895132901):0.001885076425):0.003759756
457):0.005170131503):0.000130720435,((Euphrasia:0.05859314002,Bellardia:0.009892686779):0.0
4461356787,(Orthocarpus:0.03486002774,Castilleja:0.01851342602):0.01798203312):0.008038926
367):0.003831311607):0.003901570638):0.008336249406):0.007707703756):0.02506695889,(Ehret
ia:0.04090634594,((Tournefortia:0.02570496312,Heliotropium:0.01820972354):0.03493080084,(Na
ma:0.03230674371,(Cordia:0.04195016126,(Echium:0.0274371686,(Amsinckia:0.02730054798,(Cyn
oglossum:0.02273693656,Myosotis:0.01248878636):0.01874774134):0.005473266066):0.07052046
899):0.006018981504):0.004950965978):0.03707810397):0.05315108602):0.008179147517):0.0014
93295626,((Poranopsis:0.07975446728,((((Merremia:0.005365972132,((Argyreia:0.01060491242,Ipo
moea:0.005705843542):0.008141489638,(Calystegia:0.004593322837,Convolvulus:0.007034611499
):0.01683094127):0.0007849700406):0.005491366277,Operculina:0.0271305114):0.0004791448942
,((Evolvulus:0.04596467085,Jacquemontia:0.06878862065):0.01522584251,Stictocardia:0.02041552
718):0.02783447):0.008179339958,((Dichondra:0.04438287719,Bonamia:0.02211484952):0.003895
247659,Cressa:0.05548006439):0.005542377386):0.01195658543):0.06267550328,((Nicotiana:0.00
3370428135,((Lycium:0.03975684203,Nicandra:0.005031409987):0.005175392556,(Datura:0.00476
9356272,(((Solanum:0.007125718306,Capsicum:0.05546827736):0.00514822073,Brugmansia:0.009
44061935):0.003578934018,Physalis:0.01980851876):0.003000513253):0.008639037755):0.004353
650648):0.01290382382,(Brunfelsia:0.04411281451,Cestrum:0.04808277218):0.009101306805):0.0
5153687899):0.0393263423):0.02454031628):0.01606546033,(Quintinia:0.02473713032,((Morina:0
.01951337803,(Leycesteria:0.01516128224,Lonicera:0.0294897144):0.01873210223):0.0004178409
205,(Centranthus:0.05488566582,Scabiosa:0.05305215869):0.0121851723):0.04054636951):0.0108
6446215):0.003627453233,((((((((Eryngium:0.04322032734,Sanicula:0.006873710133):0.027126988

71,(Schizeilema:0.005653803841,(Stilbocarpa:0.02153876522,Azorella:0.0394826276):0.002594088
728):0.0485992941):0.00569574116,(((Aciphylla:0.008878052166,(Cyclospermum:0.03633658536,C
ryptotaenia:0.0299714543):0.01689861396):0.02528717663,((Lilaeopsis:0.0240210337,(Chaerophyll
um:0.03063506764,(Daucus:0.01657422869,Torilis:0.006580840096):0.008348288496):0.00405095
827):0.004004568765,((Spermolepis:0.0209123474,Angelica:0.01598243003):0.005124982933,((Peu
cedanum:0.009258397316,Coriandrum:0.02057424967):0.002252619374,((Petroselinum:0.0143125
8534,(Anethum:0.00331463806,Foeniculum:0.0000010000005):0.00534564204):0.002480178689,A
pium:0.01349274922):0.009637824879):0.0000010000005):0.004777897453):0.006648557575):0.0
000010000005,Anisotome:0.01013891009):0.02884027763):0.01058888224,((Schefflera:0.0012451
50872,Panax:0.03752637536):0.007950242303,(((Pseudopanax:0.008549898141,(Osmoxylon:0.046
84190232,Griselinia:0.04327248121):0.0005086663937):0.0203849655,(Raukaua:0.007325888225,C
heirodendron:0.005030406607):0.00403287049):0.001956650174,(Hydrocotyle:0.0639005289,((Pol
yscias:0.02161497633,Meryta:0.01785582141):0.006121806114,(Centella:0.06946217139,(Fatsia:0.
003150951004,Hedera:0.0386019063):0.0000010000005):0.008852513359):0.003193883632):0.002
124247082):0.001915828656):0.003212120745):0.004992523255,Pittosporum:0.03695533755):0.0
1993959619,Sambucus:0.0354708647):0.005602576035,Pennantia:0.05105230915):0.00936280754
4,Ilex:0.06161143215):0.001853665014,(((Wahlenbergia:0.04635690674,Triodanis:0.03443159741):
0.06237891185,(Lobelia:0.03469662727,((Hippobroma:0.01240806166,Isotoma:0.03843655093):0.0
03221717381,(((Clermontia:0.01870917652,Cyanea:0.01097104984):0.005174950499,(Delissea:0.00
2760761484,Brighamia:0.005655532589):0.008538017624):0.003736135323,Trematolobelia:0.0139
178895):0.0000010000005):0.0166615292):0.0307964216):0.04488038881,((Donatia:0.0269807148
9,((Oreostylidium:0.08939845794,(Phyllachne:0.007805169845,Forstera:0.009494023784):0.017449
16762):0.039030213,Liparophyllum:0.07443303861):0.005719551336):0.005291816647,(((Selliera:0
.01309355988,Scaevola:0.007919616967):0.06305611048,(((Tussilago:0.008164551742,Haastia:0.01
063890053):0.0000010000005,(Roldana:0.01190356829,(((Crassocephalum:0.002198556291,Erechti
tes:0.0000010000005):0.0372410529,Delairea:0.01414357252):0.01123497311,(Senecio:0.0057014

49282,Emilia:0.02190896745):0.002246467168):0.01038815487):0.00873435264):0.01221006796,((
(Osteospermum:0.001707855948,Chrysanthemoides:0.001613725323):0.01200478548,Calendula:0.
01224904697):0.02043209726,(((Pleurophyllum:0.01608406611,((((Erigeron:0.00145409362,Conyza
:0.0000010000005):0.002892698323,(Solidago:0.01203289592,Heterotheca:0.01612218504):0.0081
44733126):0.0000010000005,(Brachyscome:0.005126005797,Bellis:0.0217837456):0.001559054932
):0.006377525542,Celmisia:0.001703104363):0.0000010000005):0.00317262843,(Olearia:0.012551
98017,(Ozothamnus:0.0000010000005,Pachystegia:0.01849052687):0.0000010000005):0.00539486
1075):0.01357140226,((((((Hypochaeris:0.01704783612,Picris:0.01660903023):0.006570087291,Tra
gopogon:0.02790002588):0.001654456965,Lactuca:0.02283054238):0.003216729612,(Hesperoman
nia:0.01907590913,(((Arctotheca:0.02926201019,(Symphyotrichum:0.01043868598,(Cirsium:0.0014
03440269,Carduus:0.01155047689):0.01675276069):0.01102719148):0.003114109907,((Reichardia:
0.009888668864,Sonchus:0.004614830666):0.01592445979,(Youngia:0.001861162431,(Crepis:0.036
69086678,Lapsana:0.007278207287):0.005031107579):0.003784319351):0.005416947043):0.00486
9124887,((Gerbera:0.03737927659,(Pluchea:0.03271095984,(Centratherum:0.01085797564,(Gymn
anthemum:0.01882928917,Elephantopus:0.02097792184):0.007374272095):0.004657682351):0.00
6344481712):0.01040087455,(((Pilosella:0.01994000716,(Aster:0.02846767437,Arctium:0.04284384
356):0.0000010000005):0.0173250364,(Centaurea:0.02675333593,Petasites:0.01491095286):0.004
504532786):0.005111031508,Taraxacum:0.01922129131):0.01888842198):0.006629618607):0.0092
07263051):0.002074541689):0.0146146914,(Centipeda:0.03237931975,(Sigesbeckia:0.00608114614
9,((Madia:0.0227919487,(Gaillardia:0.02574098598,(Flaveria:0.01011833776,(Tagetes:0.014977441
19,Thymophylla:0.007428343808):0.0000010000005):0.004239566072):0.002917083112):0.000001
0000005,(((((((Cosmos:0.0137350351,Bidens:0.04392072782):0.005233789519,Coreopsis:0.0149697
0049):0.0210858484,((Palafoxia:0.004970207747,Florestina:0.01049722711):0.01808940998,Acanth
ospermum:0.03681468817):0.02211740694):0.03506938484,Tridax:0.0108455657):0.01700269929,
((Perityle:0.006719390919,(Dichrocephala:0.0000010000005,Galinsoga:0.001602512339):0.0233771
0799):0.005893646934,(Montanoa:0.01431030542,Ageratina:0.01887407984):0.006771382171):0.0

01535826244):0.00134745895,(Mikania:0.01022290157,((Chromolaena:0.02767252943,(Ageratum:
0.008587674913,Adenostemma:0.0155332802):0.002029091108):0.004268722766,Eupatorium:0.0
07920284747):0.005738463315):0.006238081433):0.001293973967,(((Synedrella:0.01519017061,C
alyptocarpus:0.005983961918):0.01360643625,(Eclipta:0.003172902412,(Sphagneticola:0.01650521
767,(Wedelia:0.006985202825,(Melanthera:0.01444191123,Lipochaeta:0.002495948724):0.003197
933578):0.005866382089):0.003000247911):0.005842971483):0.007465907541,((Parthenium:0.005
972638529,(((Helianthus:0.02998675533,Zinnia:0.01829436425):0.001807187774,(Xanthium:0.0104
5623884,Ambrosia:0.009576720024):0.003019100859):0.0000010000005,Encelia:0.009830648054):
0.004334560822):0.005827959146,Verbesina:0.005961628123):0.0000010000005):0.005798994469
):0.001263849002):0.003987567509):0.009234601456):0.01701858449):0.003185628184,(Abrotane
lla:0.02748396616,((Syncarpha:0.007339959666,(Ewartia:0.003410379508,((Euchiton:0.0046886251
65,((Helichrysum:0.00666852468,Pseudognaphalium:0.001793148978):0.002101880303,Gamochaet
a:0.007157170314):0.0006510094885):0.005125065355,(Raoulia:0.008632203887,(Leucogenes:0.01
372618808,Craspedia:0.01373456522):0.0000010000005):0.0000010000005):0.001355351989):0.00
1915507855):0.01500930568,((Soliva:0.003486174924,Cotula:0.009924356472):0.031424784,((Leuc
anthemum:0.003326127068,(Artemisia:0.007131846246,Chrysanthemum:0.00173168827):0.00176
232405):0.01623546831,(Anthemis:0.01528048226,Achillea:0.007751949827):0.005992867486):0.0
3245698366):0.007392287654):0.005020814899):0.001747988041):0.004899172697):0.003504026
325):0.001321356338):0.01842450347):0.04146521484,(Alseuosmia:0.06211536627,Corokia:0.0525
4151836):0.007210563155):0.01005175626):0.00601745224):0.03851361057):0.00181133325):0.01
341543227,Merrilliodendron:0.06798200354):0.006156836824):0.01002755744):0.002106498599):
0.003706669616,Gunnera:0.06354492159):0.01844165759):0.03832489729):0.004699717153):0.01
913059732,Trimenia:0.1415319338):0.02097011799):0.1416427023):0.02095787099,Ginkgo:0.0497
2212569):0.03255016159,(Araucaria:0.1159866639,((Taxus:0.05553991115,((Cupressus:0.00928717
0078,Juniperus:0.007298798042):0.04360893019,Cryptomeria:0.0243877064):0.03922110498):0.04
054807876,Podocarpus:0.1545589763):0.01837153183):0.02344396718):0.02254137073,(Ephedra:

0.2302638687,Gnetum:0.2116068344):0.1646338371):0.03789432596,Picea:0.04160974899):0.010
73637336,(Pinus:0.04550041386,Cedrus:0.02291490972):0.005660450986):0.01310975346,Tsuga:0.
05576718872):0.01431592605,Abies:0.01431592605);

**Supplementary data 8.** Results of Phylocom analyses. Distance matrices showing floristic and ethnomedicinal distances (mean nearest-taxon distance).

**Floristic distance**

| | | | | | | | |
|---|---|---|---|---|---|---|---|
| **Hawaii** | 0 | | | | | | |
| **Northern Mariana Islands** | 0.040199 | 0 | | | | | |
| **Marquesas** | 0.038639 | 0.027131 | 0 | | | | |
| **New Zealand** | 0.048901 | 0.079826 | 0.072534 | 0 | | | |
| **Rotuma** | 0.044492 | 0.02898 | 0.029752 | 0.082209 | 0 | | |
| **Samoa** | 0.052063 | 0.034576 | 0.035099 | 0.084304 | 0.034826 | 0 | |
| **Tonga** | 0.058409 | 0.042849 | 0.049782 | 0.093356 | 0.045792 | 0.034888 | 0 |

**Ethnomedicinal distance**

| | | | | | | | |
|---|---|---|---|---|---|---|---|
| **Hawaii** | 0 | | | | | | |
| **Northern Mariana Islands** | 0.107326 | 0 | | | | | |
| **Marquesas** | 0.077634 | 0.128601 | 0 | | | | |
| **New Zealand** | 0.090217 | 0.145282 | 0.118475 | 0 | | | |
| **Rotuma** | 0.09243 | 0.120284 | 0.067679 | 0.129079 | 0 | | |
| **Samoa** | 0.080005 | 0.112426 | 0.067952 | 0.120582 | 0.063986 | 0 | |
| **Tonga** | 0.078923 | 0.115947 | 0.077837 | 0.110765 | 0.063332 | 0.05924 | 0 |
